# Engaging the complexity of diet and healthy aging in humans

**DOI:** 10.1101/2021.03.12.435077

**Authors:** Alistair M. Senior, Véronique Legault, Francis B. Lavoie, Nancy Presse, Pierrette Gaudreau, Valérie Turcot, David Raubenheimer, David G. Le Couteur, Stephen J. Simpson, Alan A. Cohen

## Abstract

Little is known about how normal variation in dietary patterns in humans affects the aging process, largely because both nutrition and the physiology of aging are highly complex and multidimensional. Here, we apply the nutritional geometry framework to data from 1560 older adults followed over four years to assess how nutrient intake patterns affect the aging process. Aging was quantified via blood biomarkers integrated to measure loss of homeostasis. Additionally, we extend nutritional geometry to 19 micronutrients. Salient results include benefits of intermediate protein and vitamin E intake. Broadly, we show that there are few simple answers of “good” or “bad” nutrients – optimal levels are generally intermediate, but dependent on other nutrients. Simpler linear/univariate analytical approaches are insufficient to capture such associations. We present an interactive tool to explore the results, and our approach presents a roadmap for future studies to explore the full complexity of the nutrition-aging landscape.

**Impact Statement:** Multidimensional nutritional analyses reveal how the association between diet and healthy aging is hard to untangle, as most nutrients have non-linear and interactive effects in humans.

## Introduction

How does what we eat affect our healthspan and longevity? The answer to this relatively concise question is unavoidably complex. Conventional approaches to understanding the effects of diet on health and aging, particularly in human nutrition, have usually focussed on single nutrients or a handful of dietary attributes/patterns (*1–3*). Yet, nutrients have both individual and interactive effects. For example, at the most macro-level, protein, carbohydrate and fat energy sources interact to determine metabolic, physiological and cognitive functioning (e.g. (*4–6*)). Similarly, the numerous phenotypic changes that occur with age are increasingly recognized as interconnected and multidimensional (*7–10*). Thus seemingly distinct physiological components of aging likely reflect a broader loss of homeostasis in a complex dynamic system rather that independent processes (*11*). Such interdependencies mean that the atomised interpretation of the effects of a single nutrient, diet, molecular mechanism or biomarker is likely to be context-dependent (*12–15*); a consequence being that the results of univariate studies are spurious and/or more difficult to reproduce, leading to inconsistent conclusions between studies.

The Geometric Framework for Nutrition (GFN) is a state-space approach to nutrition that deals with dietary complexity by considering multiple dimensions of nutrient intake simultaneously (*16, 17*). Using fully factorial experiments, the GFN has shown that components of biological aging are affected by the ratio of dietary macronutrients independently of the effects of net energy intake. These effects have been observed across taxa (*18–20*). For example, mice subjected to a lifetime of low-protein (within the boundaries of what can support growth and development), high-carbohydrate intake display improved cardiometabolic health and increased median age at death relative to animals with higher protein or fat intakes (*4, 21*). Any such benefits seemingly disappear in old-age, though, when higher protein reduces late-life mortality (*21*). These experimental findings in mice are consistent with epidemiological studies in humans that emphasise the importance of dietary protein for the elderly (reviewed in (*22, 23*)). Ecological analyses of human nations have also found that even at this highest order level, the macronutrient composition of the food supply is a powerful predictor of international variation in patterns of mortality (*24*). Collectively, these studies emphasise the importance of multi-dimensional thinking in nutrition.

The biological aging process is no more tractable than nutrition. There is no clear consensus as to what aging is (*25*), though most researchers now agree it is multi-factorial (*26, 27*). Different methods for quantifying aging correlate poorly with one another after chronological age adjustment (*28*), implying that aging is a compound process. An emerging approach is to measure the effects of aging via the breakdown in homeostatic regulation (i.e. dysregulation) across physiological systems (*10, 29*), an approach complementary to “biological age.” A statistical distance can be used to quantify how abnormal (i.e. dysregulated) an individual’s biomarker profile is, either globally or within specific systems. In this way it is possible to generate dysregulation scores that are predictive of a wide array of health outcomes during aging, including mortality, frailty, and chronic diseases (*10, 30–33*). Dysregulation scores thus simultaneously provide a proxy for the aging process and general health state. In this sense, dysregulation may be a better metric of health than measures of the aging process *per se*, if such a thing actually exists.

Here, we apply the GFN to model the effects of nutrient intake on dysregulation and aging scores in community-dwelling older adults (>67 years old). We use data from the Quebec Longitudinal Study on Nutrition and Successful Aging (NuAge) (*34*), which is extensive enough to permit application of the GFN in an epidemiological context. We show that combining these two methods provides a means to integrate the complexity inherent to nutrition and aging physiology. We begin by testing the hypothesis, suggested by the mouse data, that higher protein intakes during old age are associated with markers of improved health, while also testing whether protein interacts with the intake of the other macronutrients, notably whether an increase in the ratio of carbohydrate to protein reduces markers of aging. We then show how the same approach can be used inductively to detect micronutrient-interactions that have systemic effects.

## Results

### Overview

Briefly, NuAge participants were community-dwelling men and women, aged 67-84 years in the Montreal, Laval, or Sherbrooke areas in Quebec (Canada). They were selected randomly from the Quebec Medicare database (n=36,183), after stratification for age and sex. Individuals with good general health were recruited (n=1,793) between November 2003 and June 2005 (T1) (*34*). Participants were re-examined annually for 3 years (T2, T3 and T4). Dietary intake data were collected annually using 3 non-consecutive 24-hour diet recalls. Of the original recruits, 1,754 (98%) agreed to the integration of their data and biological specimens into the NuAge Database and Biobank for future studies. Measured from serum/plasma samples, 30 biomarkers were used to calculate dysregulation globally and for five systems that a previous study validated as being largely independent (*10*): 1) Oxygen transport; 2) Liver/kidney function; 3) Leukopoiesis; 4) Micronutrients; and 5) Lipids (see Table 1 for biomarkers in each score). We also calculated two other integrative, clinical-biomarker-based measures of biological aging: phenotypic age (PhenoAge) and Klemera-Doubal biological age (*35–37*).

**Table 1.** Biomarkers used to calculate each dysregulation score, and their population mean ± standard deviation (SD).

| Biomarker | System | Mean ± SD |
| --- | --- | --- |
| Mean corpuscular hemoglobin, MCH (pg) | Oxygen transport | 31.0 ± 1.6 |
| MCH concentration (g/L) | Oxygen transport | 340 ± 8 |
| Mean corpuscular volume (fL) | Oxygen transport | 91.3 ± 4.3 |
| Red blood cell count ($10^{12}/L$) | Oxygen transport | 4.49 ± 0.43 |
| Red cell distribution width<sup>a</sup> | Oxygen transport | 0.136 ± 0.010 |
| Hemoglobin (g/L) | Oxygen transport | 139 ± 13 |
| Albumin (g/L) | Liver/kidney function | 43.2 ± 2.4 |
| Creatinine ($\mu\text{mol}/L$)<sup>a</sup> | Liver/kidney function | 80.3 ± 21.1 |
| Albumin-globulin ratio | Liver/kidney function | 1.52 ± 0.22 |
| Bilirubin, total ($\mu\text{mol}/L$)<sup>a</sup> | Liver/kidney function | 10.2 ± 4.3 |
| Alkaline phosphatase (U/L)<sup>a</sup> | Liver/kidney function | 78.3 ± 24.5 |
| Alanine aminotransferase (U/L)<sup>a</sup> | Liver/kidney function | 12.5 ± 22.5 |
| Aspartate aminotransferase (U/L)<sup>a</sup> | Liver/kidney function | 22.9 ± 22.8 |
| γ-Glutamyltransferase (U/L)<sup>a</sup> | Liver/kidney function | 33.0 ± 32.5 |
| Lactate dehydrogenase (U/L)<sup>a</sup> | Liver/kidney function | 158 ± 30 |
| Proteins, total (g/L) | Liver/kidney function | 71.9 ± 4.1 |
| Uric acid ($\mu\text{mol}/L$) | Liver/kidney function | 342 ± 82 |
| Monocytes (differential count) | Leukopoiesis | 0.076 ± 0.029 |
| Neutrophils (differential count) | Leukopoiesis | 0.616 ± 0.087 |
| Leukocytes ($10^9/L$)<sup>a</sup> | Leukopoiesis | 6.36 ± 1.71 |
| Lymphocytes (differential count) | Leukopoiesis | 0.294 ± 0.081 |
| Folate (nmol/L)<sup>a</sup> | Micronutrients | 38.3 ± 12.5 |
| Vitamin B12 (pmol/L)<sup>a</sup> | Micronutrients | 431 ± 183 |
| β-Carotene ($\mu\text{mol}/L$)<sup>a</sup> | Micronutrients | 3.40 ± 5.55 |
| α-Tocopherol ($\mu\text{mol}/L$)<sup>a</sup> | Micronutrients | 28.6 ± 16.0 |
| γ-Tocopherol ($\mu\text{mol}/L$)<sup>a</sup> | Micronutrients | 3.36 ± 2.72 |
| Triglycerides (mmol/L)<sup>a</sup> | Lipids | 1.63 ± 0.77 |
| Cholesterol, total (mmol/L) | Lipids | 5.12 ± 1.02 |
| Cholesterol, high-density (mmol/L) | Lipids | 1.44 ± 0.39 |
| Platelet count ($10^9/L$)<sup>a</sup> | — | 237 ± 64 |

<sup>a</sup> Variable was log-transformed to meet assumptions of normality.

We assessed the effects of intakes of macronutrients and micronutrients, as well as their interactions, on measures of dysregulation and aging. Our primary tool was the generalised additive model (GAM), a form of multiple regression that tests for non-linear multidimensional effects using ‘smooth’ terms (*38, 39*). We explored three-dimensional effects of nutrient intake. Because GAMs can estimate non-linear effects, qualitative interpretation of the sign of estimated effects comes through visualisation (rather than a single regression coefficient). Here effects were visualised using the nutrient intake surfaces common to the GFN. Importantly, GAMs can be used to correct for factors (e.g. sociodemographic status) that might be expected to confound relationships, in the same way as regression might be used in epidemiology. For each outcome we fitted a series of eight models with varying levels of correction (factors explored were income, education level, age, physical activity, number of comorbidities, sex and current smoking status; see text S1).

The main text contains a complete cases analysis comprising 3569 observations from 1560 people. In the supplementary materials (text S2 and S3) we report sensitivity analyses, where we have imputed missing income data, and also analysed a more exclusive subset of the complete cases dataset (see Table S1 for population summaries; the exclusive dataset excluded diabetics, individuals on prescribed diets, BMI <22 or >29.9, or with substantial weight fluctuation). These sensitivity analyses estimated similar effects to those in the main text. However, for the exclusive dataset, in many places these effects (although qualitatively similar) do not meet the criteria for statistical significance. We interpret this latter point as evidence that the effects in the two datasets are similar, but that the power of the complete cases analysis is required to detect statistical significance.

### Dietary Macronutrients

Our first model, model 1, tested for the effects of macronutrient intake (in kJ/day) on outcomes without any statistical corrections (see text S1 for parameters of all models). We detected statistically significant effects of macronutrient intake on liver/kidney and micronutrient system dysregulation scores, as well as biological age (Figures 1a through 1c; see supplementary materials Table S2 for all statistical model output). Model 1 predicted a relatively minor effect of protein on liver/kidney function dysregulation, and non-linear effects of all carbohydrates and fats on liver/kidney function dysregulation. Individuals who consumed high (> 6000 kJ/day) or low (< 3000 kJ/day) levels of carbohydrates typically had slightly elevated (around 0.25 SD above the population mean) dysregulation scores (Figure 1a). Very high intakes of lipids (> 4000 kJ/day; note this is within 2SD above the mean lipid intake) had the highest liver/kidney function dysregulation (∼0.4 SD above the mean; figure 1a). With regard to micronutrient dysregulation scores, individuals with moderately high carbohydrate (5000 to 6000 kJ/day) and low lipid (< 2000 kJ/day) intake had minimal dysregulation. Again, high lipid intake was associated with maximal dysregulation (Figure 1b). Subjects with protein intake around 2000 kJ/day were predicted by model 1 to have low dysregulation (Figure 1b). Biological age was predicted to be the lowest for individuals with high levels of intake for all three macronutrients (Figure 1c); note that in this analysis nutrient intakes are not expressed relative to any measure of individual requirements.

**Figure 1.**
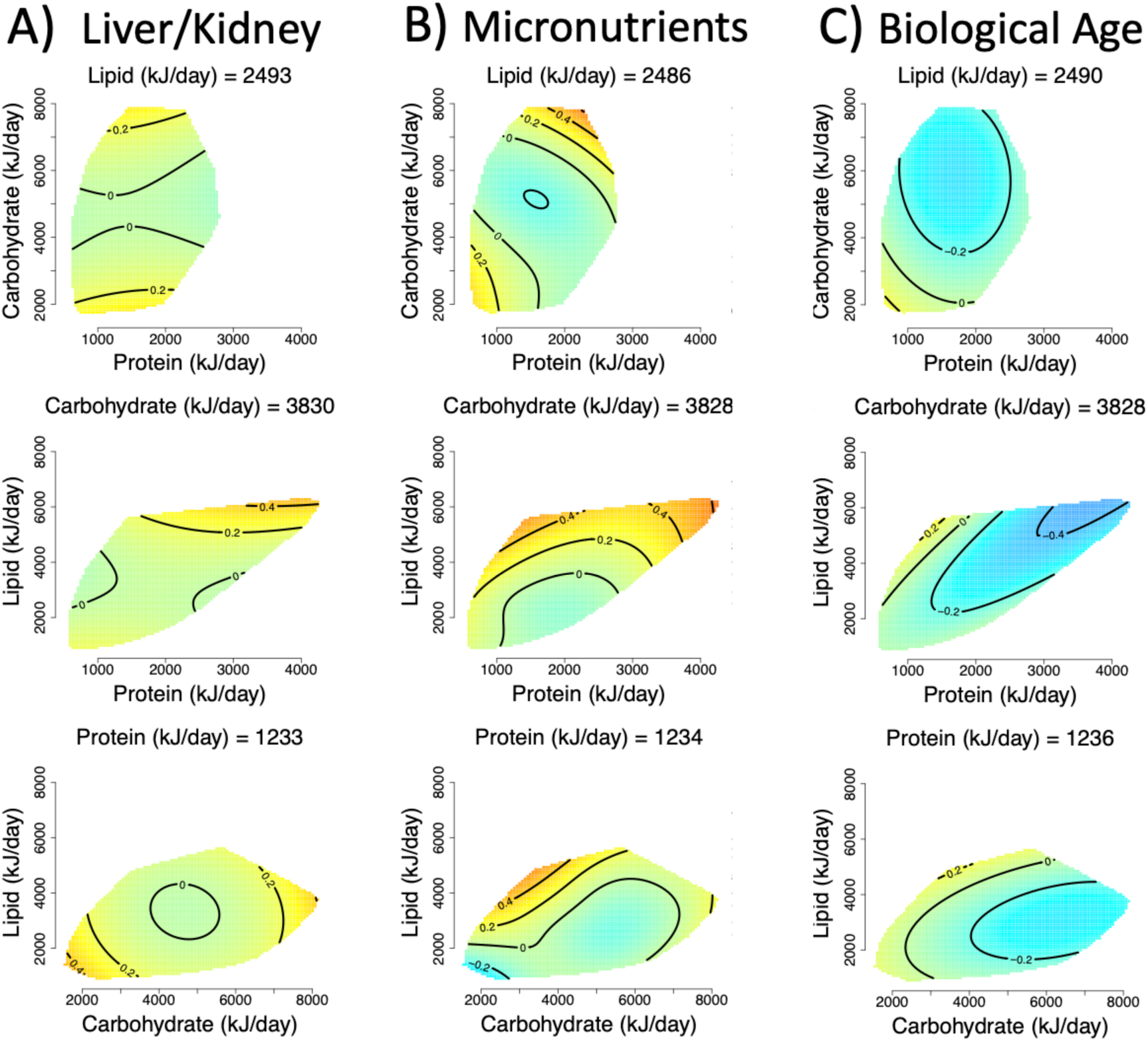
Effects of total dietary intake (kJ/d) of protein, carbohydrates, and fats on A) liver/kidney function dysregulation (GAM three-way smooth term: edf=9, Ref. df=9, F=2.66, p<0.01, Dev. Expl.=1.5%), B) micronutrient dysregulation (GAM three-way smooth term: edf=14.6, Ref. df=18.1, F=1.8, p<0.05, Dev. Expl.=2.21%) and C) biological age score as predicted by model 1 (GAM three-way smooth term: edf=9, Ref. df=9, F=4.06, p<0.001, Dev. Expl.=2.79%). Surfaces across the top row show effects of protein (x-axis), and carbohydrate (y-axis) intake, those across the middle row protein and lipid, and the bottom row is carbohydrate and lipid. The third macronutrient is held at the values given on all panels (population median). Warm colours indicate high dysregulation, and cool colours low dysregulation. All scores were Z-transformed to one SD, and surfaces colours are scaled such that deep blue and red represent effects of at least -0.8 and 0.8 (conventionally considered an effect of large biological magnitude (*83*)).

In model 2 we considered dysregulation score as a function of macronutrient intake relative to what we define as the typical energy intake given height, weight, age, sex and physical activity (text S4). We again detected effects of macronutrient intake on liver/kidney function and micronutrients dysregulation and biological age score, but in addition, effects on PhenoAge were detected (Figures 2a through 2d; supplementary materials Table S3). Again, this analysis indicated high protein intake relative to what is typical (100% above average) had low liver/kidney function dysregulation scores (Figure 2a). Interestingly, both PhenoAge and biological age were predicted to be minimised at elevated carbohydrate levels and typical levels of lipid and protein (Figures 1c and 1d). These effects remained after making corrections for potential confounders, both excluding and including number of comorbidities (models 3 and 4; supplementary materials, Tables S4 and S5), and showed a similar trend within the exclusive dataset (see text S3 and Figure S3).

**Figure 2.**
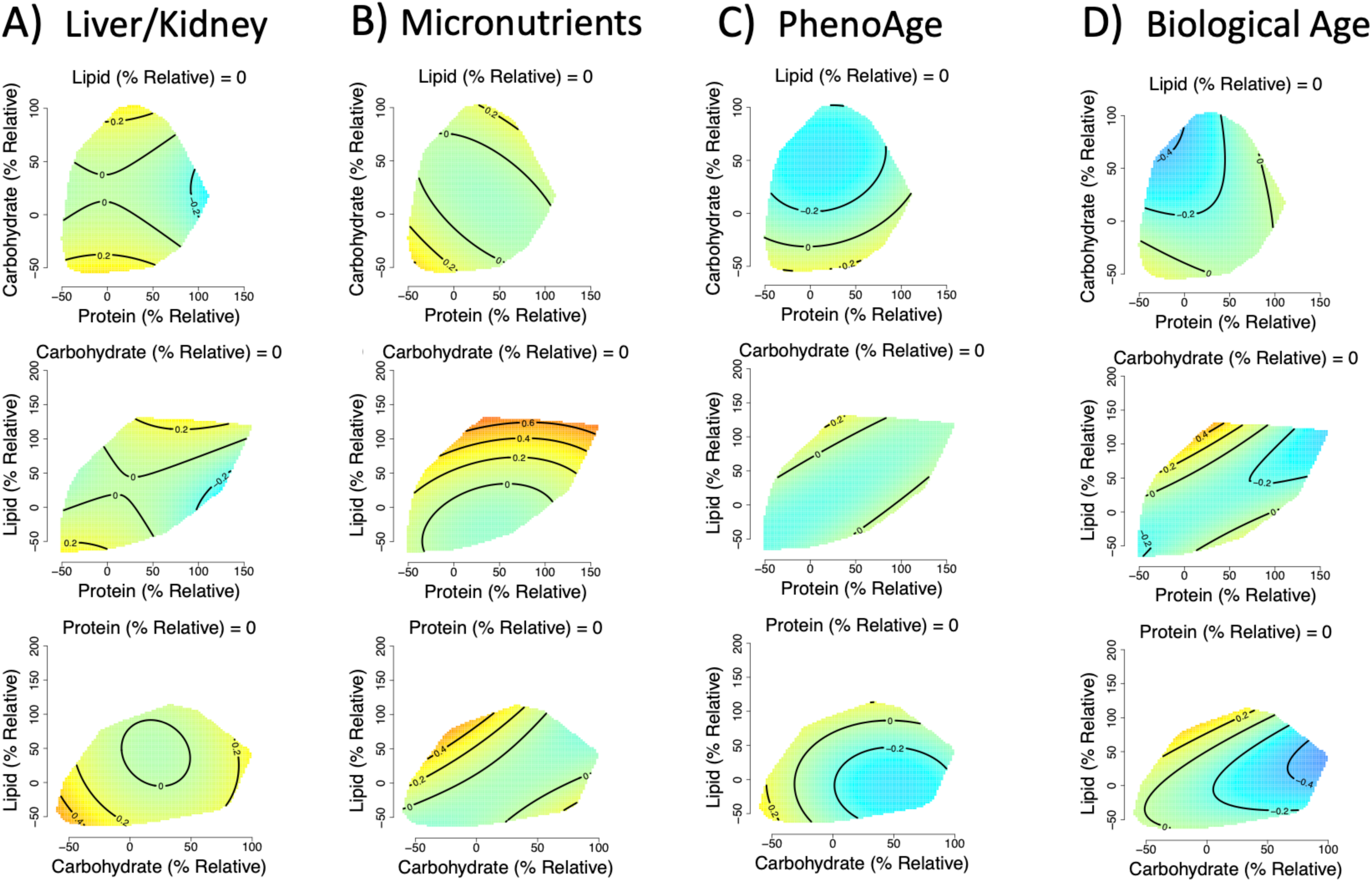
Effects of relative dietary macronutrient intake (relative to the required intake based on age, weight, height, sex and physical activity level; see supplementary materials, text S4) on A) liver/kidney function dysregulation, (GAM three-way smooth term: edf=9, Ref. df=9, F=3.8, p<0.001, Dev. Expl.=1.8%), B) micronutrient dysregulation, (GAM three-way smooth term: edf=9, Ref. df=9, F=2, p<0.05, Dev. Expl.=0.9%), C) PhenoAge (GAM three-way smooth term: edf=9, Ref. df=9, F=2, p<0.05, Dev. Expl.=0.9%) and D) biological age (GAM three-way smooth term: edf=9, Ref. df=9, F=2, p<0.05, Dev. Expl.=0.9%) score as predicted by model 2. Surfaces across the top row show effects of protein (x-axis), and carbohydrate (y-axis) intake, those across the middle row protein and lipid, and the bottom row is carbohydrate and lipid. The third macronutrient is held at the values given on all panels. Warm colours indicate high dysregulation, and cool colours low dysregulation. All scores were Z-transformed to one SD, and surfaces colours are scaled such that deep blue and red represent effects of at least -0.8 and 0.8 (conventionally considered an effect of large biological magnitude (*83*)). Individuals with a relative intake value of 100, eat 100% more of that macronutrient per day (in kJ) than is predicted to be typical for the population given their age, sex, weight, height and level of physical activity level. Conversely, individuals with a relative intake value of 0 would eat the required amount of that macronutrient per day.

### Dietary Micronutrients

Intakes of a number of the micronutrients explored were strongly correlated (figure 3). Hierarchical clustering based on correlation distance suggested seven clusters of micronutrients with very highly correlated intakes (*r* > 0.65). The first principal component (PC1) of intakes within each of the seven clusters explained between 90 and 100% of the variation of intake of micronutrients within that cluster (supplementary materials Table S6). PC1 estimates uniformly displayed positive correlations with intakes of micronutrients within the clusters. For subsequent analyses, cluster-specific PC1 values were used as measures of intakes of micronutrients within clusters.

**Figure 3.**
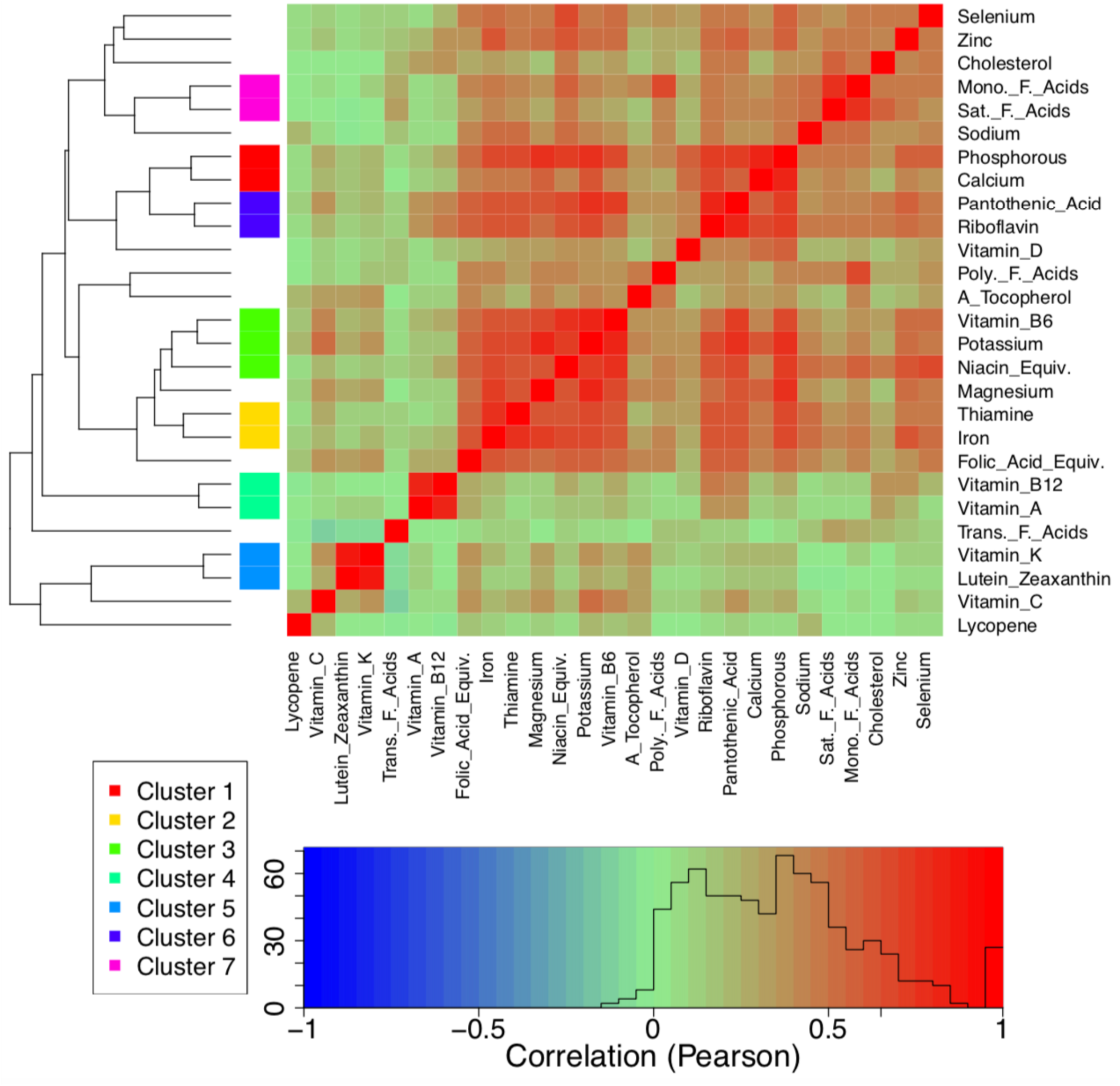
Correlogram of the strength of correlations (Pearson’s correlation) between intakes of micronutrients. Correlations have been clustered hierarchically based on correlation distance (dendrogram). On the basis of this clustering, we grouped micronutrients with highly correlated intakes (> 0.65) for subsequent dimension reduction using principle components analysis.

After using PCA to reduce micronutrient dimensionality, we were left with 19 minimally correlated micronutrient variables (PC1 of clusters or individual micronutrients), making it feasible to run GAMs for all 969 3-way micronutrient combinations for each dysregulation score (micronutrient-specific models; see supplementary materials text S1). For all scores except lipid dysregulation there were a greater number of significant micronutrient smooth terms than would be expected under the null hypothesis (Figures 4a through 4h). After correction for the false discovery rate (FDR), there were no significant effects of micronutrient intake on oxygen transport or lipid dysregulation scores. There were 905 models that detected significant effects for at least one score, although 363 of these were solely related to aging scores (i.e., no significant effect on any dysregulation score; supplementary materials Table S7). There were 17 combinations for which we detected significant effects on all scores (except oxygen transport and lipid dysregulation). Interestingly all 17 models contained *α*-tocopherol (vitamin E) as one of the three micronutrients. Arguably any one of these micronutrient combinations may be of interest and could warrant further investigation in response to *a priori* hypotheses (note results from all models detecting significant interactions can be found at http://nuage.cohenlab.ca). Here however we focus on the effects of *α*-tocopherol, vitamin C and trans-fatty acids because this combination had the highest mean percent deviance explained across all scores.

**Figure 4.**
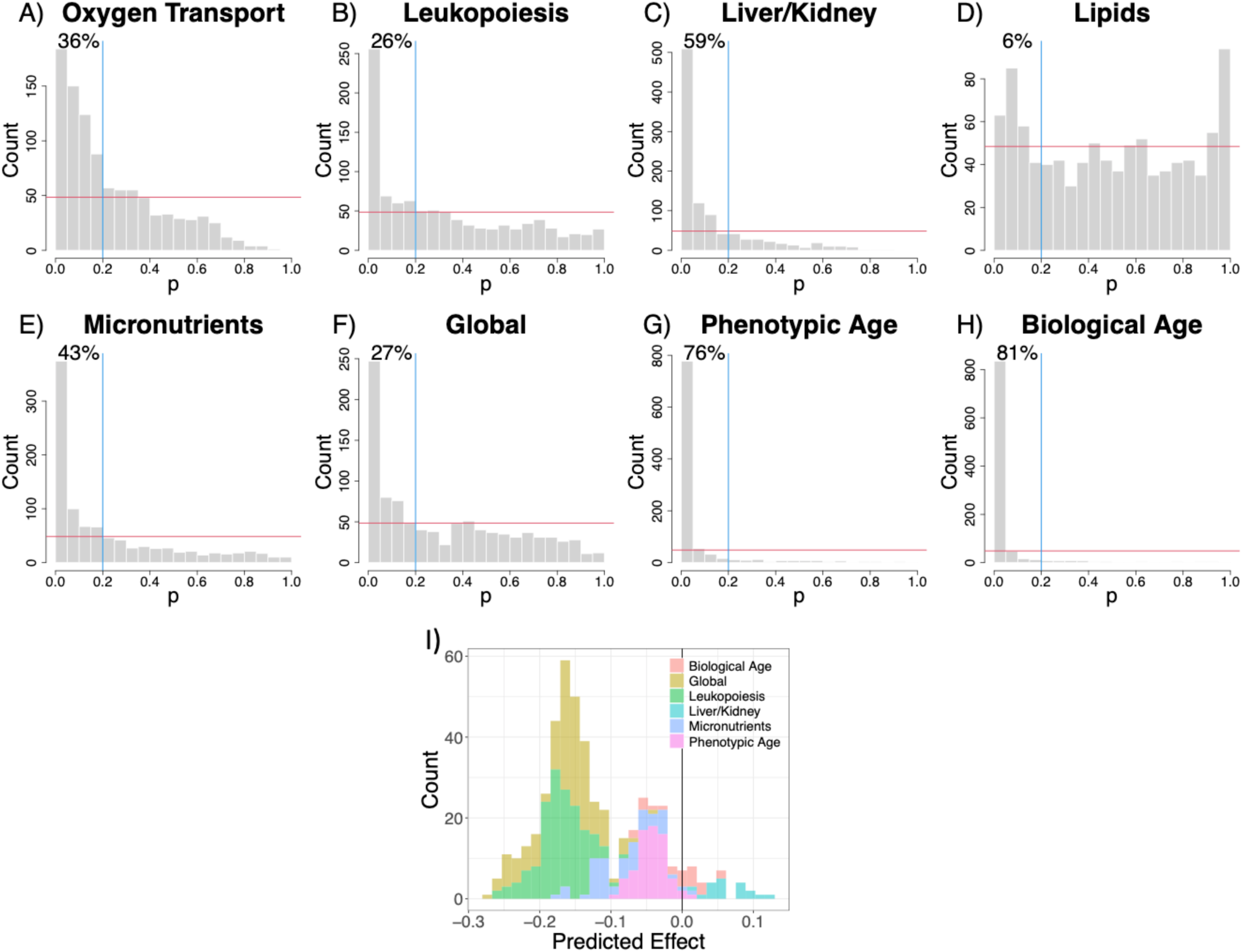
Frequency histograms for the p-values for smooth terms for the 969 unique three-way combinations of micronutrient intakes as given by GAMs (micronutrient-specific models; see supplementary materials, text S1) where A) oxygen transport, B) leukopoiesis, C) liver/kidney function, D) lipids, E) micronutrients, F) global dysregulation, G) PhenoAge and H) biological age score was treated as an outcome. The red horizontal line indicates the expected frequency under the null hypothesis (that the outcome is unaffected by micronutrient intake) and the blue vertical line demarks p=0.2. The percentage of p-values falling into the upper left quadrant is given. I) Frequency histogram of the effects of an increase in *α*-tocopherol intake of 2 SD from the population average as predicted by different models. Predictions come from all models containing significant three-way micronutrient smooth terms involving *α*-tocopherol, and make adjustments as per model 6. Predictions assume population average values for all other intakes (including alcohol), income, education level, age, physical activity level (PASE), number of comorbidities, men and non-smoker.

In models 5 and 6 we tested for effects of these three micronutrients with correction for confounders, excluding and including comorbidities respectively. In these models we detected effects of micronutrients on leukopoiesis, liver/kidney function, micronutrients and global dysregulation (supplementary materials Tables S8 and S9). The effects varied slightly across the four systems. (Figures 5a through 5d for model 6). For example, elevated intakes of trans-fatty acids are predicted to be detrimental for liver/kidney dysregulation, but have a minor beneficial effect on leukopoiesis dysregulation. Nevertheless, consuming around 2SD of *α*-tocopherol above the population mean, while consuming vitamin C and trans-fatty acids at the population mean results in low dysregulation across the four scores, suggesting a systemic benefit of high, but not excessive, *α*-tocopherol intake (Figure 5). To ensure that increased *α*-tocopherol does not have detrimental interactions with other micronutrients, we screened the complete list of significant micronutrient combinations (after FDR adjustment) for those that included *α*-tocopherol intake; 153 combinations were identified for at least one outcome score. Figure 4I shows the distribution of dysregulation scores (all traits) predicted for a 2SD increase in *α*-tocopherol for any identified effects of micronutrients (after inclusion for potential confounders). For all scores, intakes of *α*-tocopherol 2SD above our sample mean (mean ± SD = 4.75 ± 2.73) is predicted to lead to reductions in dysregulation or aging *Z*-scores (below the population average) or changes close to 0 (the population average).

**Figure 5.**
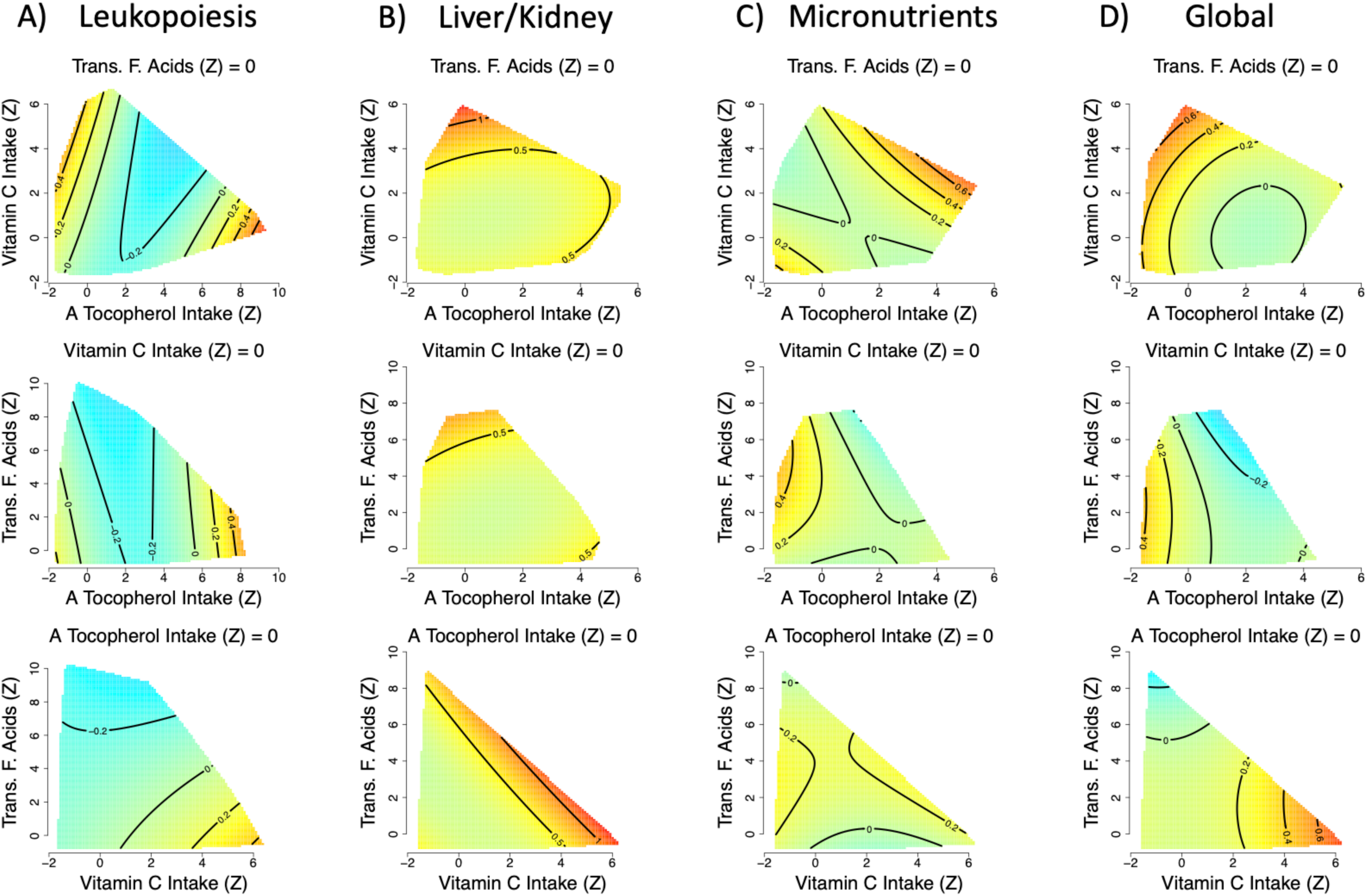
Effects of total dietary micronutrient (*α*-tocopherol, vitamin C and trans-fatty acid intake) intake on A) leukopoiesis (GAM three-way smooth term: edf=9, Ref. df=9, F=3.8, p<0.001, Dev. Expl.=6.7%), B) liver/kidney function (GAM three-way smooth term: edf=9, Ref. df=9, F=1.9, p<0.05, Dev. Expl.=3.7%), C) micronutrients (GAM three-way smooth term: edf=9, Ref. df=9, F=2.3, p=0.01, Dev. Expl.=3.8%) and D) global (GAM three-way smooth term: edf=9.2, Ref. df=9.5, F=2.5, p<0.01, Dev. Expl.=7.5%) dysregulation score as predicted by model 6. Intakes have been *Z*-transformed and are thus in units of SD. In all cases predictions assume the micronutrient not displayed on either the x- or y-axis is held at the population mean. Numeric confounding variables included in model 6 were alcohol intake, income, education level, age, physical activity level (PASE) and number of comorbidities, and predictions assume population mean values. Predictions are for men and assume a non-smoker.

### Macro, Micro or Both

We found statistical support for effects of both macro and micronutrients on liver/kidney function and micronutrient dysregulation. There can be strong covariances among macro and micronutrient intake owing to their co-occurrences in foods (see supplementary materials text S5). For example, vitamin C intake is often considered to be a marker of fruit and vegetable consumption (*40*), and unsurprisingly in this dataset high vitamin C intake coincides with a high carbohydrate diet (text S5). One may thus question which of these two nutritional levels is a better predictor of dysregulation, or if both must be considered. We evaluated this by fitting the two three-dimensional effects for macro and micronutrients simultaneously (models 7 and 8, excluding and including comorbidities respectively) and assessing model fit relative to corresponding previous models using Akaike information criterion (AIC). For liver/kidney function dysregulation a model including macronutrients is favoured by AIC (Table 2). In contrast, for the micronutrient dysregulation score a model including only micronutrients is favoured by AIC (Table 2). We also note here that even the most complex models fitted, which include both macro and micronutrients, as well as other potential predictors of health status, explain less than 5% of the deviance in these dysregulation scores for this population (Table 2).

**Table 2.** Akaike information criterion (AIC) and deviance explained (%) by models 4, 6, and 8 for liver/kidney and micronutrients dysregulation. These models contained sex, smoking status, alcohol intake, income, education level, age, physical activity (PASE) and number of comorbidities, alongside the nutritional predictors stated. See supplementary materials Table S10 for model variants excluding number of comorbidities.

| Dysregulation Score | Model Number | Nutrient Predictors | AIC | Deviance Explained |
| --- | --- | --- | --- | --- |
| Liver/kidney function | Model 4 | Macronutrients | 5172 | 4.1% |
|  | Model 6 | Micronutrients | 5179 | 3.69% |
|  | Model 8 | Macro + Micronutrients | 5177 | 4.82% |
| Micronutrients | Model 4 | Macronutrients | 4946 | 3.63% |
|  | Model 6 | Micronutrients | 4943 | 3.81% |
|  | Model 8 | Macro + Micronutrients | 4945 | 4.66% |

## Discussion

Here we used multidimensional modelling techniques to test for effects of nutrient intake on physiological dysregulation in an aged population. We find that consuming above average protein and *α*-tocopherol intake, which is the most active form of ‘vitamin E’, is associated with lower levels of physiological dysregulation. In contrast, above average carbohydrates (given what is typical for height, weight, age sex and physical activity) coupled with typical protein and lipid intakes minimised measures of biological aging.

Analysis of the effects of dietary macronutrients on age-specific mortality in mice has shown that protein restriction when coupled with increased carbohydrate intake, extends median lifespan through reduced mortality in middle life, but that higher protein in late life may reduce mortality (*21*). Given that our study population consisted entirely of older adults, our findings with regards physiological dysregulation are consistent with this experimental literature. Our results are also concordant with numerous studies highlighting the need for increased protein intake in older people, in particular to offset sarcopenia and decreased physical performance associated with aging (reviewed in (*22, 23*)). It is interesting to speculate, though, whether reapplying our methods to middle aged cohorts would detect similar benefits of elevated protein intake; the experimental literature, and some data in humans, suggests perhaps not (*20, 21, 41*).

Numerous micronutrient interactions have been demonstrated in the experimental absorption literature (e.g., calcium-iron, vitamin C-iron etc. (*42*)). Our analysis indicates that such interactions may have strong enough effects on health in old age to be detected at the level of the population. For example, elevated vitamin E intake can produce either low or high micronutrient dysregulation scores, depending upon the level of vitamin C intake (Figure 5C). *In vitro* and *in vivo* studies, including a supplementation trial in healthy adults, have detected vitamin C-E absorption interactions, including where simultaneous supplementation leads to elevated circulating levels, and where vitamin C can reduce oxidised vitamin-E recovering its role as an antioxidant (*43–45*). Thus, there is a mechanistic basis for what we have detected, and it is certainly worth testing for interactions between these micronutrients in other epidemiological cohorts.

We found that consuming *α*-tocopherol at 2SD above the population average is associated with benefit; in the subject population this level corresponds to 10.21mg/day of *α*-tocopherol. The World Health Organisation recommended intake of *α*-tocopherol for those aged 65+ is 7.5mg/day in females and 10mg/day in males, thus the value we highlight is not beyond current guidelines (*46*). Highlighting the importance of considering non-linearity, more extreme intake patterns (e.g., >4 SDs above average) were associated with harm (Fig. 5). This finding accords with the results of RCTs suggesting excessive vitamin E supplementation may increase all-cause mortality (*47*). We also detected a non-linear effect of carbohydrate intake on dysregulation, which suggests individuals at the upper/lower extremes of the observed carbohydrate intakes suffer poor health. Epidemiological and meta-analytic study of the effects of carbohydrates on all-cause mortality in humans has found identical patterns (*48*). Generally speaking, our study therefore provides further support to the importance of looking beyond ‘single nutrient at a time’ and monotonic “more is better” analyses (*1, 49*), to detect interactive effects of nutrients. For space reasons we have not presented and discussed the full range of results for all micronutrient intake interactions detected here. However, we have illustrated an approach that can be used to study the effects of multiple micronutrient intakes on health. The GAM-based approach can either be used for discovery (i.e., as means to detect micronutrient interactions), or as a targeted mechanism-driven paradigm to test specific hypotheses about micronutrient interactions derived from experimental biology or nutrition science (*42, 50*). Additionally, we encourage readers to explore http://nuage.cohenlab.ca where the effects of micronutrient intakes on dysregulation and aging scores in this population can be visualized interactively, and *a priori* hypotheses explored.

Even in our most complex models, the deviance explained remains at around 5%. This is within the bounds of what analyses of other outcomes in this cohort have found (e.g. (*51*)). Nevertheless, the surfaces we present here show relatively large effects of nutrition on dysregulation levels, which is potentially important as nutrition is readily modifiable. The low deviance explained may be due to a number of factors including: the reporting errors, and systematic reporting biases inherent in epidemiological studies of nutrition (*52–54*); the fact we consider additive effects of macro and micronutrients rather than interactions (more complex models are theoretically possible but are limited by sample size and visualisation beyond three-dimensions); and the rarity of the extreme dietary profiles at the edges of the surfaces, where the strongest effects are found (the standard error of the surface is a proxy for sample density; e.g. supplementary materials figure S6). An implication of this latter point is that some of the largest effects of nutrient intake we report will only apply to a small proportion of the population, and that our physiology is often robust enough to tolerate relatively wide variation without much consequence. Similar patterns are observed when using the GFN to map evolutionary fitness to organisms that ecologists pre-define as dietary ‘generalists’ (*16, 55–57*). This is consistent with an understanding of nutrition in which our ancestors evolved to tolerate an array of dietary patterns (*58*). Accordingly, homeostasis can be maintained across a wide array of nutritional states, with the caveat that when diet becomes too extreme dysregulation can increase very rapidly (“falling off a dietary cliff”). The tolerance for different diets – the size of the plateau in this analogy – could of course vary as a function of genetic or environmental factors that predispose us to greater risk (*59*).

In contrast to dysregulation, we found that PhenoAge and biological age were minimised on relatively high carbohydrate and lower to moderate protein intakes. Similar results have been observed in numerous experimental studies on the effects of protein to carbohydrate ratio on aging in model organisms (*20*). The somewhat discordant results between dysregulation measures and PhenoAge / Biological Age are not necessarily that surprising. Increasingly, the literature shows that aging is multivariate and heterogeneous (*10, 28*). In this context, our results imply that different dietary patterns come with their own benefits and drawbacks in their effects on the different facets of aging. A practical consequence of this specificity is that dietary recommendations could be tailored to slow the most advanced aging processes based on an individual’s biological profile. Of course, replication in other cohorts is needed to confirm the precise patterns we report, and even then, context-specific clinical recommendations will be essential.

An important next step for analyses of this nature will be to complement them with analyses of the effects of food/broad dietary patterns on physiological dysregulation (*60*). The intakes of individual nutrients are often markers for broader dietary patterns. High vitamin C/E intake can reflect a diet rich fresh fruit and vegetables (*40*), whereas a high trans-fatty acid intake may reflect a diet comprised of processed foods. Such analyses may help to elucidate why, for example, we detect a minor beneficial effect of trans-fatty acids on leukopoiesis dysregulation, where other studies have detected the opposite (*61*); it is possible that in this population high trans-fatty acid intakes represent a diet high in dairy, as opposed to processed foods.

Previously, studies have applied GFN thinking to experimental and observational contexts in humans. They have shown that the macronutrient composition of the diet is associated with biomarkers of cardiovascular health, energy intake, obesity and specific chronic diseases (*62–67*). Important as they are, these analyses were concerned with unidimensional measures of health and healthspan. Here, for the first time we have used the GFN to model holistic measures of systemic functioning during aging. How results from experimental nutrition and geroscience map on to humans living in the community remains unclear. In part, our understanding has been hampered by a lack of techniques that cut across the complexity inherent to nutrition and biological aging in real-world contexts; the approach we present here is a promising start. Future applications could include personalized approaches to aid in healthy aging, and screening of at-risk older adults to ensure they do not fall off the “dietary cliff.” Finally, our results advocate against the popular practice of eating to maximize or minimize certain nutrients. The dose-response relationship is often U-shaped, and highly dependent on context (e.g., age, other aspects of diet); targeting in the absence of clear evidence is likely to do more harm than good.

## Methods

### Participant Information and Dietary Intake Data

The NuAge Database and Biobank and the present study have been approved by the Research Ethics Board (REB) of the CIUSSS-de-l’Estrie-CHUS (Quebec, Canada; projects 2019-2832 and 2015-868/14-141, respectively). The original sample includes 1587 individuals recruited between November 2003 and June 2005 (T1) to which 206 volunteers were added (*34*). They were community-dwelling men and women, aged 67-84 years, able to speak English or French, and in good general health at recruitment. Notably, they had to be free of disabilities in activities of daily living, not cognitively impaired and able to walk 300 metres or to climb 10 stairs without rest. A structured interview was conducted annually in the NuAge Study at baseline (T1) and for the next 3 years (T2, T3 and T4) to gather the following data. Sociodemographic (actual income, education) and lifestyle (smoking, alcohol) information was obtained using a general study questionnaire developed from standard health survey questions (*68*). The number of self-reported chronic health conditions (i.e. comorbidities) was computed from an adaptation of the Older Americans Resources and Services (OARS) Multidimensional Functional Assessment questionnaire (*69*). Chronic health conditions considered were self-reported cancer (within the last 5 years); hypertension; self-reported liver or gallbladder disease; self-reported surgery of the digestive system; self-reported heart trouble; self-reported circulation trouble in arms or legs; self-reported thrombosis, cerebral haemorrhage, or cerebrovascular accident; self-reported transient cognitive impairment; self-reported Parkinson disease; self-reported diabetes; self-reported emphysema or chronic bronchitis; self-reported asthma; rheumatoid arthritis, arthritis, or rheumatism; self-reported osteoporosis; self-reported kidney diseases; self-reported thyroid and gland problems; self-reported surgery of the circulatory system; and mini-mental state examination (MMSE) total score between 22 and 26. Usual physical activity was assessed using the Physical Activity Scale for the Elderly (PASE) questionnaire, with higher scores corresponding to higher physical activity level (*70*). Weight and height were measured, and body mass index (BMI = weight [kg] / height [m]^2^) calculated for each participant (*34*).

Dietary intake data were collected annually (T1 to T4) using 3 non-consecutive 24-hour diet recalls (*71–73*). Each set included 2 weekdays and 1 weekend day, with the first administered during the annual face-to-face interview and the others by telephone interviews without prior notice. Based on the USDA 5-step multiple-pass method (*74*), interviewers recorded a detailed description and portion sizes of all foods and beverages consumed by each participant the day before the interview. Only energy and nutrients coming from foods were considered (i.e., excluding supplements). All interviewers were trained registered dietitians. Energy and nutrient intake were computed from the 24-hour diet recalls using the CANDAT-Nutrient Calculation System (version 10, ©Godin London Inc.) based on the 2007b version of the Canadian Nutrient File (CNF) from Health Canada and a database of 1,200 additional foods that was developed on site (*75*).

### Biomarker data and Dysregulation Score

Biomarkers were chosen based on our previous study (*76, 77*), their clinical use, and according to their availability in NuAge. In total 30 biomarkers were used to calculate dysregulation globally and for five systems that a previous study validated as being largely independent (*10*): 1) Oxygen transport; 2) Liver/kidney function; 3) Leukopoiesis; 4) Micronutrients; and 5) Lipids (Table 1).

Not all individuals had complete data on biomarkers and nutrition, so sample size depended on the physiological system and the analysis. For the oxygen transport and leukopoiesis systems, sample size varied between 1224 and 1331. For the other systems, the sample size varied between 654 and 730. The difference in sample size is due to a more extensive serum biomarker analysis conducted in 2016 on a subsample of ∼750 individuals that were selected at random among the 904 of the 1754 participants who met the following criteria: (1) the individual needed to have blood sampling conducted without any missing intermediate visits; (2) the individual needed at least two visits with blood samples; and (3) there had to be a sufficient number of stored aliquots at all visits to analyze the selected biomarkers.

Dysregulation was calculated using the Mahalanobis distance (D_M_), a statistical distance which measures how deviant or aberrant an individual’s biomarker profile is compared to a reference population. Using D_M_, a global dysregulation score (“Global”) was calculated with all 30 biomarkers and by physiological system for each participant at each visit, as previously described (*76*). Higher scores indicate higher dysregulation. Age at recruitment in NuAge is restricted from 67 to 84 years, thus, to construct our reference population we used data from the first visit, which represents a population in general good health, according to the inclusion and exclusion criteria (22). Some biomarkers were log- or square-root-transformed as necessary to approach normality. All biomarkers were then centred at the mean of the reference population and divided by the standard deviation of the reference population to standardize them before calculation of dysregulation scores.

Alongside the 6 measures of physiological dysregulation, we also included two other measures of biological aging: phenotypic age (PhenoAge) and Klemera-Doubal biological age. PhenoAge measures the predicted age of the individual based on the individual’s mortality risk as assessed via 9 biomarkers and calibrated in NHANES IV. Biological age measures the age of an individual as predicted based on linear projections of a set of age-associated biomarkers. We calculated PhenoAge as described elsewhere (*36, 37*), although because CRP (C-reactive protein) is not available in the NuAge cohort, we used the mean CRP value from NHANES IV 1999-2010 (*37*) for all individuals. PhenoAge showed a weaker correlation with chronological age in our population than previously reported (*r*=0.67 *vs* 0.96; (*37*)). Exploratory analysis of other datasets in our possession indicated the discrepancy was likely due to the limited age range rather than the absence of CRP (data not shown). Calculation of biological age was based on previous work by Levine (*35*). We first searched for biomarkers that correlated with chronological age and found 13 with *r* > 0.1 (*p* ≤ 0.05). After removing three biomarkers with numerous missing values (non-HDL cholesterol, LDL, and estimated glomerular filtration rate), we were left with the following list: hemoglobin, hematocrit, red blood cell count, red cell distribution width, monocyte count, albumin, folate, creatinine, blood urea nitrogen, and lymphocyte percentage. Using these biomarkers, we calculated biological age as previously described (*35, 78*).

### Analyses

All analyses were performed in the statistical programming environment R (*79*). In all cases dysregulation scores were first log transformed, and then *Z*-transformed (mean centred and divided by one SD) prior to model fitting. Analysis was performed using GAMs (*38, 80*). GAMs are a form of the generalised linear (mixed) models (GLMs, and GLMMs) that allow the user to model complex non-linear effects through the inclusion of non-parametric ‘smooth’ terms (usually implemented as a spline). These terms can be specified alongside the conventional linear parametric terms in a linear model. Smooth terms can be additive in a single dimension, or multi-dimensional to test for synergistic interactions. The statistical significance of smooth terms can be interpreted via a p-value, but their effect must be interpreted visually (as opposed to numerically via a regression coefficient). GAMs were implemented using the ‘gam’ function in the R package *mgcv*, and terms estimated by restricted maximum likelihood (*38, 39*).

We first tested the hypothesis that increased protein is associated with improved health in old age. We estimated the effects of daily intake of macronutrients (protein, carbohydrate and fat, in kJ) on each dysregulation/aging score using GAMs. In all models, dysregulation/aging score at an observation was the outcome. In model 1, macronutrient intakes at those observations were fitted as a three-dimensional smooth term (thin-plate spline), and individual subject ID was fitted as a random effect (there are multiple observations per individual). However, the dataset is made up of individuals whose energy requirements likely vary due to height, weight, age, sex and physical activity level. Thus, we also estimated macronutrient intakes relative to what is typical for an individual in this population given the aforementioned variables (estimated from the residuals of a model of intake; see supplementary materials text S4). In model 2, we modelled dysregulation scores as a function of relative macronutrient intake using a GAM as above.

In model 3, we tested for the effects of confounders by refitting model 2 to include potential confounders as additive effects; sex (men / women), smoking status (current / not current), current income, number of years of education, alcohol intake (g/d), physical activity level (PASE) and age were included. We ran a separate model with comorbidities as an additional potential confounder (model 4). Numeric predictors were *Z*-transformed prior to fitting and were included in the model as smooth terms, and categorical predictors as parametric terms.

In the second part of our analyses, we considered the effects of those micronutrients for which intake data were available. Micronutrient intakes are likely to be highly correlated. Therefore, we performed hierarchical clustering on correlations of micronutrient intakes based on correlation distance using the ‘hclust’ function in *base* R. For any clusters of highly correlated micronutrients we performed a principal component analysis (PCA; ‘prcomp’ function in *base* R) and used the first principal component (PC1) as our measure of intake for the nutrients within that cluster. The number of micronutrients adds complexity to understating their interactive effects because: 1) the estimation of multidimensional smooth terms in GAMs becomes challenging as the number of dimensions grows, and 2) the smooth terms in GAMs must be interpreted visually. For these reasons we restricted ourselves to considering 3 micronutrient dimensions at a time. For each three-way combination of micronutrient intake (PC1 for clusters) we fitted a GAM with a three-dimensional smooth term for the three micronutrients and a random effect for individual. We then identified those models with significant effects after correction for the false discovery rate (FDR, *q* < 0.05; (*81*)). We note that correction by FDR assumes that p-values are independent, which is not the case here. However, overlooking this non-independence is conservative in that it will result in fewer significant effects rather than more, and thus we proceeded without correction. In the main text we interpret specific cases of interest, however, all results can be accessed at http://nuage.cohenlab.ca. We also tested for effects of micronutrients alongside correction for the potential confounders discussed above (models 5 and 6), and by modelling micro and macronutrient intakes simultaneously (models 7 and 8). Note that including two three-dimensional smooth terms (one for micronutrients and one for macronutrients) is not the same as a six-dimensional smooth term; this approach is substantially less power-hungry but still allows us to adjust micronutrient models for macronutrient intake, and vice-versa. See supplementary materials (text S1) for a concise list of all models implemented.

Statistical significance was inferred for terms in GAMs based on *α* of 0.05. Where we found it necessary to compare amongst models containing different predictors we used Akaike Information Criterion (AIC), where smaller values (beyond a margin of two points) indicate a better model fit (*82*). Smooth terms from GAMs were interpreted from model predictions. For three dimensional effects of nutrient intake we present two-dimensional surface plots, where intakes of nutrients are given on the x- and y-axes, while the intake of the third nutrient is held constant at a value stated on the figure panel. On all surfaces deep blue/red areas indicate low/high *Z* bio-scores (i.e., good/bad health), respectively.

The main text contains a complete cases analysis where we report results from analyses on all observations for which relevant predictor and dysregulation score data were available (i.e. inclusive dataset with missing income data excluded). Supplementary texts S2 and S3 contain the results of two sets of sensitivity analyses. In the first we have imputed missing income data (31%) as the participant-specific mean value to increase our sample size. In the second we have analysed a subset of the data (<50% of the total data) where we have excluded any observation at which the subject was recorded as either; diabetic (type-1 or 2), reported as being on a medically prescribed diet, having a BMI outside the range of 22 to 29.9, or that came from a subject with a coefficient of variation of weight > 0.04 over the course of all the observations (i.e. for whom weight fluctuated substantially).

Key analyses were performed by two separate members of the team, using broadly the same approach but with no consultation on the detailed analytical decisions (i.e. what might be inferred from seeing the Results but not the Methods). Discrepancies leading to qualitative changes in the conclusions were flagged and resolved. VL performed this validation, with AMS as primary data analyst. Datasets containing all participants’ variables used in this study and those from dietary intakes estimated from 24-hour dietary recalls were transmitted by the NuAge team in November-December 2018 and October 2019, respectively. Data for some biomarkers were obtained in a previous secondary project (*77*).

## Acknowledgements

AMS was supported by a discovery early career researcher award from the Australian Research Council (ARC DECRA: DE180101520). The NuAge cohort was supported by the Canadian Institutes of Health Research (CIHR) grant #62842. The NuAge Database and Biobank are supported by the *Fonds de recherche du Québec* (*FRQ*) grant #2020-VICO-279753, the Quebec Network for Research on Aging, a thematic network funded by the *FRQ - Santé (FRQ-S)* and by the Merck-Frosst Chair funded by *La Fondation de l’Université de Sherbrooke*. This work was supported by Canadian Institutes of Health Research (CIHR, grant #153011). AAC is supported by a CIHR New Investigator Salary Award and *FRQ-S* Senior Fellowship, and is a member of the *FRQ-S* funded *Centre de recherche du CHUS* and *Centre de recherche sur le vieillissement*. NP is a Junior 1 Research Scholar of the *FRQ-S*.

## Text S1: Models

For each dysregulation score (*Z*-transformed) we implemented GAMs as follows.

Model 1.

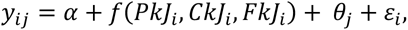

where, *y_ij_* is the *i*th dysregulation score for the *j*th individual, *α* is the intercept value, *f*(*PkJ_i_*, *CkJ_i,_ FkJ_i_*) is a three-dimensional smooth term for the effects of absolute intake of protein, carbohydrate and fat in kilojoules at observation *i*, *θ_j_* is a random effect for the *j*th individual and *ε_i_* is the residual for the *i*th observation; *θ_j_* and *ε_i_* are assumed to be normally distributed with means of 0 and SDs estimated from the data.

Model 2.

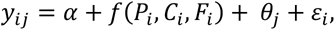

which is as in model 1, but the three-dimensional macronutrient smooth term is now relative intake of each macronutrient at the *i*th observation (estimated as described in text S4 ‘Estimation of Relative Intake’).

Model 3.

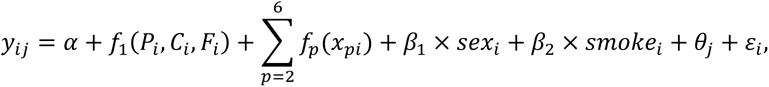

which is as in model 2 but with additional additive smoothed terms for potential confounding variables, where *f_p_*(*x_p_*) is a smoothing function for the *p*th predictor *x* (*x*_2_ = income, *x*_3_ = alcohol intake, *x*_4_ = age, and *x*_5_ = PASE, and *x*_6_ = number of years of education), at observation i. These numeric confounders were *Z*-transformed prior to model fitting. The model also includes parametric terms (*β*_1_ and *β*_2_) for the effect of sex (0 for men and 1 for women) and current smoking status at observation *i* (0 for non-smoker, 1 for smoker).

Model 4.

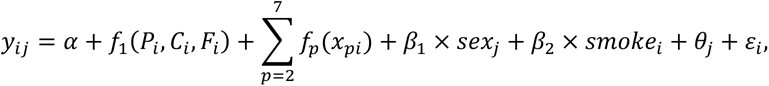

which is as in model 3 but includes a further smooth term, *x*_7_, which is number of comorbidities of the individual at observation *i*.

Micronutrient-specific models.

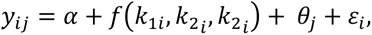

A model was specified for each unique 3-way combination of micronutrients, where *k*_1*i*_, *k*_2*i*_ and *k*_3*i*_ are the intakes (*Z*-transformed) of the *k*th combination of 3 micronutrients at the *i*th observation, and other terms are as above.

Model 5.

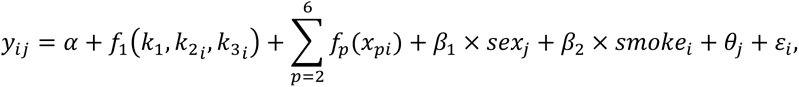

which is as in model 3 but considers a combination of three micronutrients (or PC1 of clusters thereof), *f*(*k*_1*i*_, *k*_2*i*_, *k*_3*i*_), determined to be of interest following the fitting of micronutrient-specific models.

Model 6.

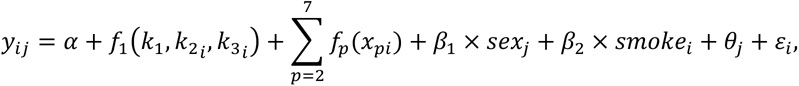

which is as in model 6 but includes the seventh smoothed term for number of comorbidities (*x*_7_).

Model 7.

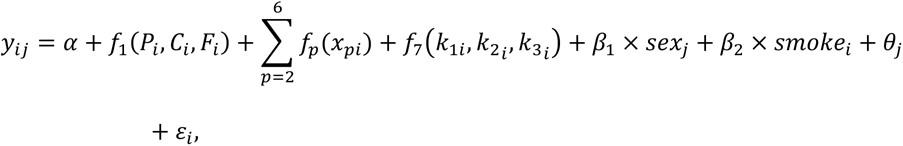

which is as in model 3 but with *f*_7_(*k*_1*i*_, *k*_2*i*_, *k*_3*i*_), which is an additional three-dimensional smooth term for the intake of a combination of three micronutrients (or PC1 of clusters thereof) determined to be of interest following the fitting of micronutrient-specific models.

Model 8.

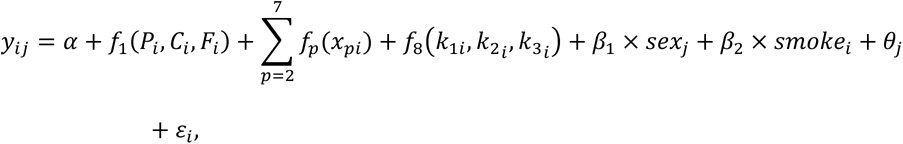

which is as in model 7 but includes the seventh smoothed term for number of comorbidities (*x*_7_).

## Text S2: Analysis of Inclusive Dataset with Imputation of Missing Income Data

Applying model 1 to the imputed dataset detected significant effects of absolute macronutrient intake on liver/kidney function dysregulation, micronutrient dysregulation and biological age. These are same outcomes as are described in the analysis in the main text, and the estimated effects of nutrition were near identical. Also, as in the main text, we detected effects of relative nutrient intake on liver/kidney function dysregulation, micronutrient dysregulation and PhenoAge and biological age scores in models 2 through 4. Figure S1 is the equivalent to figure 2 in the main text, but with the imputed income data included; the estimated effects are qualitatively identical.

We re-assessed the effects of *α*-tocopherol (vitamin E), vitamin C and trans-fatty acids on dysregulation scores in the imputed dataset using model 6 (i.e. with full correction for socioeconomic factors and comorbidities). As in the main text, we detected effects of this micronutrient combination on leukopoiesis and micronutrient dysregulation (figure S2). We did not detect effects on liver/kidney function or global dysregulation, although in the latter case the smooth term was close to reaching statistical significance (GAM three-way smooth term: edf=9, Ref. df=9, F=1.8, p=0.06, Dev. Expl.=6%). We did however detect effects of micronutrients on lipid system dysregulation (figure S2). These analyses show that lipid dysregulation is low for individuals with relatively high trans-fatty acid and low vitamin C intake; while this is an unexpected observation, this is an extreme dietary profile and the surface in this area is associated with a large degree of sampling error. Nonetheless, as in the main text, elevated levels of *α*-tocopherol (vitamin E) intake to around 2 SD are generally predicted to be beneficial in that they lead to low dysregulation scores (figure S2).

## Text S3: Analysis of Exclusive Dataset

We re-ran our analyses on a subset of the data in which we had excluded any observations where the individual was recorded as any of the following: diabetic (type-1 or 2), reported as being on a medically prescribed diet, having a BMI outside the range of 22 to 29.9, or came from a subject that had a coefficient of variation of weight > 0.04 over the course of all the observations within the dataset. The logic being that, in doing so we would be ensuring that our results were not driven by individuals who had a dietary associated chronic disease and extreme dietary profile that was disproportionately affecting our models. Table S1 shows the profiles of the populations captured within the different datasets. These restrictions excluded over the half of the data, which substantially reduced statistical power. With this reduced dataset our analyses estimated effects with the same sign as those in the main text, but for macronutrients (and liver/kidney function dysregulation with micronutrients) failed to reach the level of statistical significance.

Figure S3 displays the same results as figure 2 in the main text. For this restricted dataset the models estimated the same effects as those for the inclusive dataset, although the effects for liver/kidney function dysregulation, micronutrient dysregulation and biological age score were non-significant. Figure S4 shows the effects of *α*-tocopherol, vitamin C and trans-fatty acid intake on leukopoiesis, liver/kidney function, micronutrient and global dysregulation score. Here all potential confounding variables have been taken into account (i.e. model 6). We see the qualitatively similar effects to those presented in the main text, although the effects are non-significant for all cases but global dysregulation score.

## Text S4: Estimation of Relative Intake

To estimate relative macronutrient intake based on individual requirements, we modelled the mean daily intake of each macronutrient (protein, carbohydrate and fat, in kJ) at an observation, as a function of the subject’s weight, height, age, sex and physical activity level at that observation. Intakes of the three macronutrients were specified as outcomes in a multi-response generalised additive model (GAM), where sex was a categorical parametric predictor, weight, height, and age were fitted as a 3-dimensional smooth term (smoothed by sex), and physical activity level (PASE score) as an additional additive smoothed term. This model estimates the typical intake of each energy-yielding macronutrient as a function of these factors, which are commonly expected to affect energy requirements. From the model we estimated an individual’s relative macronutrient intake as the ratio of their residual for observed intake to that predicted by the model multiplied by 100. Thus, individuals with a relative intake value of 100, eat 100% more of that macronutrient per day (in kJ) than is predicted to be typical for this population given their age, sex, weight, height and level of physical activity. Conversely, individuals with a relative intake value of 0 eat what is the expected given these factors weight, height age, sex and PASE score.

## Text S5: Dietary Macronutrient Composition and Micronutrient Intake

We tested for associations between the composition of the diet in terms of percentage energy from the three macronutrients, protein, carbohydrates and lipids and intake of the micronutrients (or PC1 for clusters) considered. We used mixture models (MMs), also known as Scheffe’s polynomials (1), where micronutrient intake (*z*-transformed to one SD) was the outcome and percentage energy from each macronutrient were the predictors. For each micronutrient we fitted 5 MMs; MM 1 was a null model and MMs 2 through 5 tests for increasingly complex linear through non-linear effects of diet composition on micronutrient intake (see equations 1 through 4 in (1)). It is notable that MM 2 is identical to the substitution models commonly used in nutritional epidemiology (2, 3). We compared among MMs using Akaike Information Criterion (AIC; (4)), assuming that the simplest model within 2 AIC points of the lowest AIC observed was best supported. MMs were implemented using the *mixexp* package in R (1). For micronutrients of interest, we visualised the predictions for the AIC-supported MM using right-angle mixture triangles (RMTs; (5)). For all micronutrients AIC favoured models other than the null model (Table S11), suggesting that dietary macronutrient composition is associated with intake of micronutrients. Figure S5 shows the association between diet composition and the three micronutrient intakes of interest in the main text as predicted by the AIC favoured MMs; *α*-tocopherol, vitamin C and trans-fatty acids. Diets with a low percentage energy from protein and carbohydrates and high percentage energy from fat had high intake of *α*-tocopherol and trans-fatty acids (figures S5a and S5c). A high carbohydrate diet was associated with high levels of vitamin C, perhaps reflecting intake of fruits and vegetables (figure S5b).

## Supplementary Tables

**Table S1:** Summary of the variants of the dataset used (see main text). The inclusive dataset comprised 3569 observations (all time points) of 1560 individuals with missing income values excluded and 5152 observations of 1561 individuals including income values imputed. The exclusive dataset comprised 1473 observations of 567 individuals with missing income values excluded. Values shown outside of brackets are the mean and SD of observations excluding observations with missing incomes, while those in brackets correspond to those when imputed incomes are included. For dysregulation scores values are log Mahalanobis distance. For dysregulation and aging scores sample sizes vary among scores with exact values given with corresponding analyses. Note for smoking, the proportion excludes former smokers.

| Variable (units) | Inclusive Dataset (inc. imputed income) |  | Exclusive Dataset |  |
| --- | --- | --- | --- | --- |
|  | Mean | SD | Mean | SD |
| Age (years) | 76.77 (76.75) | 4.28 (4.30) | 76.68 | 4.29 |
| Income (CAD/yr) | 39514 (39473) | 22211 (22205) | 41716 | 22489 |
| Education (years) | 11.70 (11.71) | 4.52 (4.51) | 12.34 | 4.67 |
| Alcohol Intake (g/d) | 6.61 (6.50) | 11.63 (11.60) | 7.43 | 11.30 |
| Physical activity (PASE score) | 99.70 (97.45) | 51.84 (51.46) | 108.95 | 52.80 |
| Comorbidities (n) | 2.64 (2.59) | 1.66 (1.63) | 2.29 | 1.54 |
| Body weight (kg) | 72.99 (72.74) | 14.24 (14.07) | 69.51 | 9.44 |
| Height (m) | 1.62 (1.62) | 0.09 (0.09) | 1.62 | 0.09 |
| Men (proportion) | 0.5 (0.51) | NA | 1.47 | 0.50 |
| Current Smoker (proportion) | 0.06 (0.06) | NA | 0.04 | 0.20 |
| Protein Intake (kJ/d) | 1241 (1237) | 375 (376) | 1258 | 380 |
| Carbohydrate Intake (kJ/d) | 3879 (3876) | 1138 (1150) | 4017 | 1150 |
| Lipid Intake (kJ/d) | 2566 (2565) | 933 (931) | 2621 | 925 |
| α-tocopherol Intake (mg/day) | 4.75 (4.62) | 2.73 (2.73) | 4.75 | 2.76 |
| Vitamin C Intake (mg/day) | 114.8 (115.5) | 66.5 (67.1) | 118.2 | 65.9 |
| Trans-Fatty Acid Intake (g/day) | 0.83 (0.82) | 1.00 (1.00) | 0.84 | 1.01 |
| Oxygen transport Dysregulation (score) | 1.37 (1.40) | 1.02 (1.01) | 1.25 | 1.02 |
| Leukopoiesis Dysregulation (score) | 1.20 (1.08) | 1.05 (1.00) | 1.27 | 1.06 |
| Liver/kidney function Dysregulation (score) | 3.60 (3.60) | 1.00 (1.00) | 3.51 | 0.94 |
| Lipid Dysregulation (score) | 0.80 (0.76) | 0.98 (1.00) | 0.76 | 0.97 |
| Micronutrient Dysregulation (score) | 1.88 (1.90) | 1.02 (1.01) | 1.81 | 1.03 |
| Global Dysregulation (score) | 7.01 (6.91) | 1.03 (0.99) | 6.94 | 0.98 |
| PhenoAge (score) | 73.06 (72.90) | 7.29 (7.16) | 72.30 | 6.90 |
| Biological Age (score) | 76.39 (76.35) | 8.39 (8.40) | 76.12 | 8.33 |

**Table S2:** Output of model 1 applied to the inclusive data set with missing income data excluded. For parametric terms estimates, standard error (SE), t and p-values are given. For smooth terms, s(…), estimated degrees of freedom (edf), reference degrees of freedom (Ref. df), F and p-values are given. The sample size (n) is the number of observations, and the random effect for subject ID was fitted as the smooth term s(sujetno). The p-values of significant smooth terms for macronutrients are highlighted in bold.

| Dysregulation Score | Parameter | Estimate/edf | SE/Ref. df | t/F | p |
| --- | --- | --- | --- | --- | --- |
| Oxygen transport | Intercept | 0.118 | 0.034 | 3.461 | 0.001 |
| n = 3332: % Dev. Explained = 0.69: AIC = 9456 | s(PROTEIN_kJ.d,CARBO_kJ.d,LIPID_kJ.d) | 9.014 | 9.028 | 0.736 | 0.677 |
|  | s(sujetno) | 0.942 | 1.000 | 16.130 | <0.001 |
| Leukopoiesis | (Intercept) | -0.181 | 0.034 | -5.259 | <0.001 |
| n = 3334: % Dev. Explained = 2.24: AIC = 9438 | s(PROTEIN_kJ.d,CARBO_kJ.d,LIPID_kJ.d) | 20.653 | 27.031 | 0.814 | 0.745 |
|  | s(sujetno) | 0.974 | 1.000 | 36.897 | <0.001 |
| Liver/kidney function | (Intercept) | 0.082 | 0.046 | 1.779 | 0.076 |
| n = 1834: % Dev. Explained = 1.5: AIC = 5200 | s(PROTEIN_kJ.d,CARBO_kJ.d,LIPID_kJ.d) | 9.004 | 9.007 | 2.659 | **0.005** |
|  | s(sujetno) | 0.809 | 1.000 | 4.226 | 0.022 |
| Lipid | (Intercept) | -0.127 | 0.047 | -2.716 | 0.007 |
| n = 1991: % Dev. Explained = 0.98: AIC = 5654 | s(PROTEIN_kJ.d,CARBO_kJ.d,LIPID_kJ.d) | 9.013 | 9.025 | 0.738 | 0.676 |
|  | s(sujetno) | 0.905 | 1.000 | 9.559 | 0.001 |
| Micronutrient | (Intercept) | <0.001 | 0.024 | -0.002 | 0.998 |
| n = 1750: % Dev. Explained = 2.21: AIC = 4962 | s(PROTEIN_kJ.d,CARBO_kJ.d,LIPID_kJ.d) | 14.588 | 18.099 | 1.763 | **0.025** |
|  | s(sujetno) | 0.003 | 1.000 | <0.001 | 0.693 |
| Global | (Intercept) | <0.001 | 0.024 | <0.001 | 1.000 |
| n = 1718: % Dev. Explained = 1.24: AIC = 4889 | s(PROTEIN_kJ.d,CARBO_kJ.d,LIPID_kJ.d) | 13.114 | 15.842 | 0.983 | 0.473 |
|  | s(sujetno) | 0.001 | 1.000 | <0.001 | 0.959 |
| PhenoAge | (Intercept) | -0.151 | 0.049 | -3.096 | 0.002 |
| n = 1834: % Dev. Explained = 1.57: AIC = 5199 | s(PROTEIN_kJ.d,CARBO_kJ.d,LIPID_kJ.d) | 9.007 | 9.015 | 1.711 | 0.081 |
|  | s(sujetno) | 0.925 | 1.000 | 12.399 | <0.001 |
| Biological Age | (Intercept) | -0.152 | 0.049 | -3.082 | **0.002** |
| n = 1796: % Dev. Explained = 2.79: AIC = 5069 | s(PROTEIN_kJ.d,CARBO_kJ.d,LIPID_kJ.d) | 9.010 | 9.020 | 4.060 | <0.001 |
|  | s(sujetno) | 0.925 | 1.000 | 12.260 | <0.001 |

**Table S3:** Output of model 2 applied to the inclusive data set with missing income data excluded. For parametric terms estimates, standard error (SE), t and p-values are given. For smooth terms, s(…), estimated degrees of freedom (edf), reference degrees of freedom (Ref. df), F and p-values are given. The sample size (n) is the number of observations, and the random effect for subject ID was fitted as the smooth term s(sujetno). The p-values of significant smooth terms for macronutrients are highlighted in bold.

| Dysregulation Score | Parameter | Estimate/edf | SE/Ref. df | t/F | p |
| --- | --- | --- | --- | --- | --- |
| Oxygen transport | Intercept | 0.115 | 0.034 | 3.378 | 0.001 |
| n = 3332: % Dev. Explained = 1.14: AIC = 9454 | s(PROTEIN_kJ.d,CARBO_kJ.d,LIPID_kJ.d) | 13.059 | 15.951 | 1.015 | 0.419 |
|  | s(sujetno) | 0.939 | 1.000 | 15.357 | <0.001 |
| Leukopoiesis | (Intercept) | -0.183 | 0.034 | -5.320 | <0.001 |
| n = 3334: % Dev. Explained = 1.96: AIC = 9446 | s(PROTEIN_kJ.d,CARBO_kJ.d,LIPID_kJ.d) | 17.789 | 23.276 | 0.619 | 0.921 |
|  | s(sujetno) | 0.974 | 1.000 | 37.742 | <0.001 |
| Liver/kidney function | (Intercept) | 0.081 | 0.046 | 1.756 | 0.079 |
| n = 1834: % Dev. Explained = 1.8: AIC = 5194 | s(PROTEIN_kJ.d,CARBO_kJ.d,LIPID_kJ.d) | 9.004 | 9.007 | 3.298 | **0.001** |
|  | s(sujetno) | 0.805 | 1.000 | 4.120 | 0.024 |
| Lipid | (Intercept) | -0.126 | 0.047 | -2.693 | 0.007 |
| n = 1991: % Dev. Explained = 1.08: AIC = 5652 | s(PROTEIN_kJ.d,CARBO_kJ.d,LIPID_kJ.d) | 9.012 | 9.023 | 0.971 | 0.462 |
|  | s(sujetno) | 0.904 | 1.000 | 9.391 | 0.001 |
| Micronutrient | (Intercept) | <0.001 | 0.024 | <0.001 | 1.000 |
| n = 1750: % Dev. Explained = 1.09: AIC = 4968 | s(PROTEIN_kJ.d,CARBO_kJ.d,LIPID_kJ.d) | 9.012 | 9.025 | 2.116 | **0.025** |
|  | s(sujetno) | <0.001 | 1.000 | <0.001 | 0.749 |
| Global | (Intercept) | <0.001 | 0.024 | <0.001 | 1.000 |
| n = 1718: % Dev. Explained = 0.76: AIC = 4887 | s(PROTEIN_kJ.d,CARBO_kJ.d,LIPID_kJ.d) | 9.924 | 10.728 | 1.088 | 0.373 |
|  | s(sujetno) | <0.001 | 1.000 | <0.001 | 0.958 |
| PhenoAge | (Intercept) | -0.140 | 0.049 | -2.888 | 0.004 |
| n = 1834: % Dev. Explained = 1.99: AIC = 5191 | s(PROTEIN_kJ.d,CARBO_kJ.d,LIPID_kJ.d) | 9.003 | 9.006 | 2.601 | **0.006** |
|  | s(sujetno) | 0.915 | 1.000 | 10.804 | 0.001 |
| Biological Age | (Intercept) | -0.150 | 0.050 | -3.028 | 0.003 |
| n = 1796: % Dev. Explained = 1.88: AIC = 5086 | s(PROTEIN_kJ.d,CARBO_kJ.d,LIPID_kJ.d) | 9.002 | 9.005 | 2.198 | **0.020** |
|  | s(sujetno) | 0.922 | 1.000 | 11.826 | <0.001 |

**Table S4:**
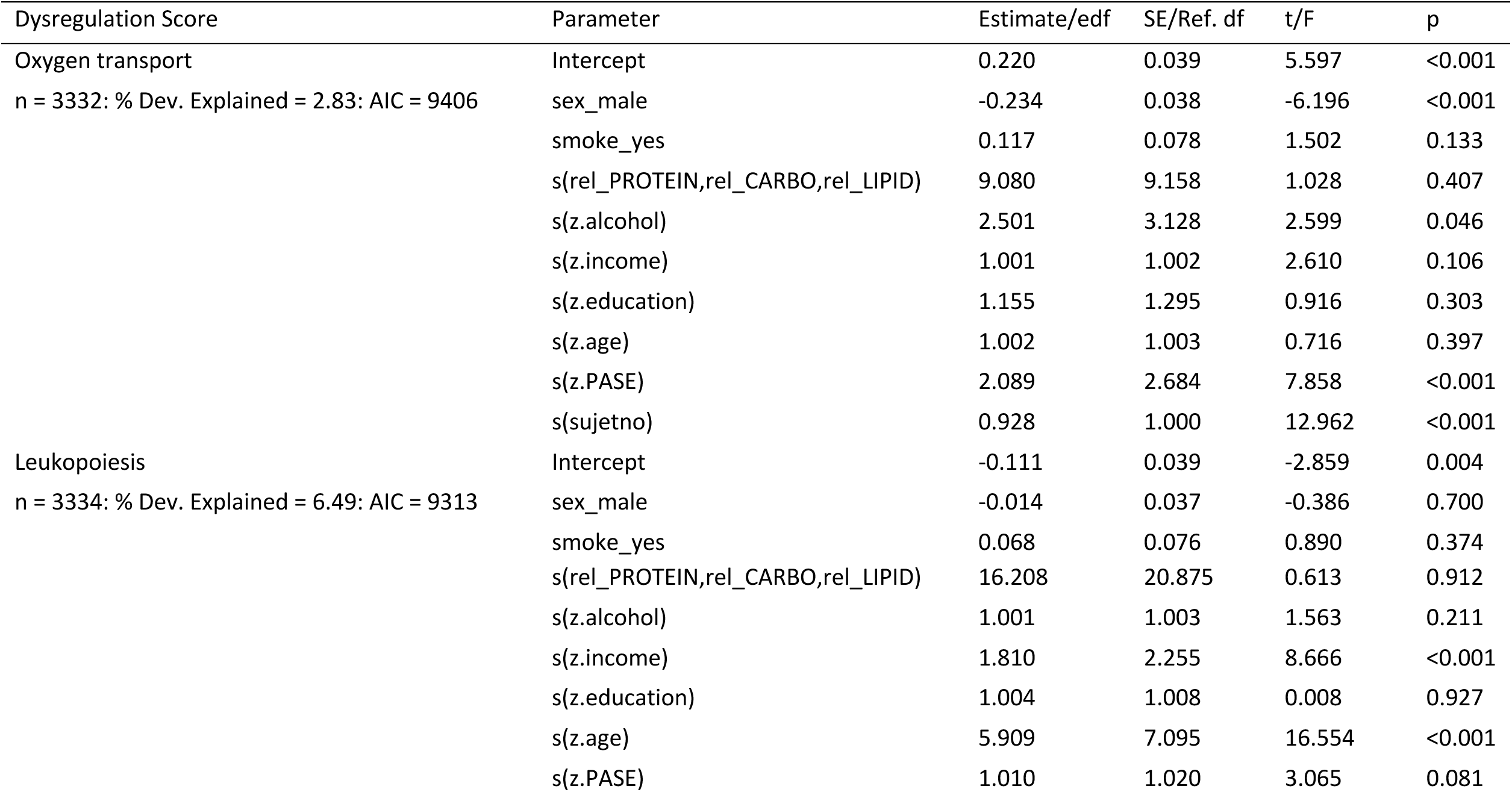
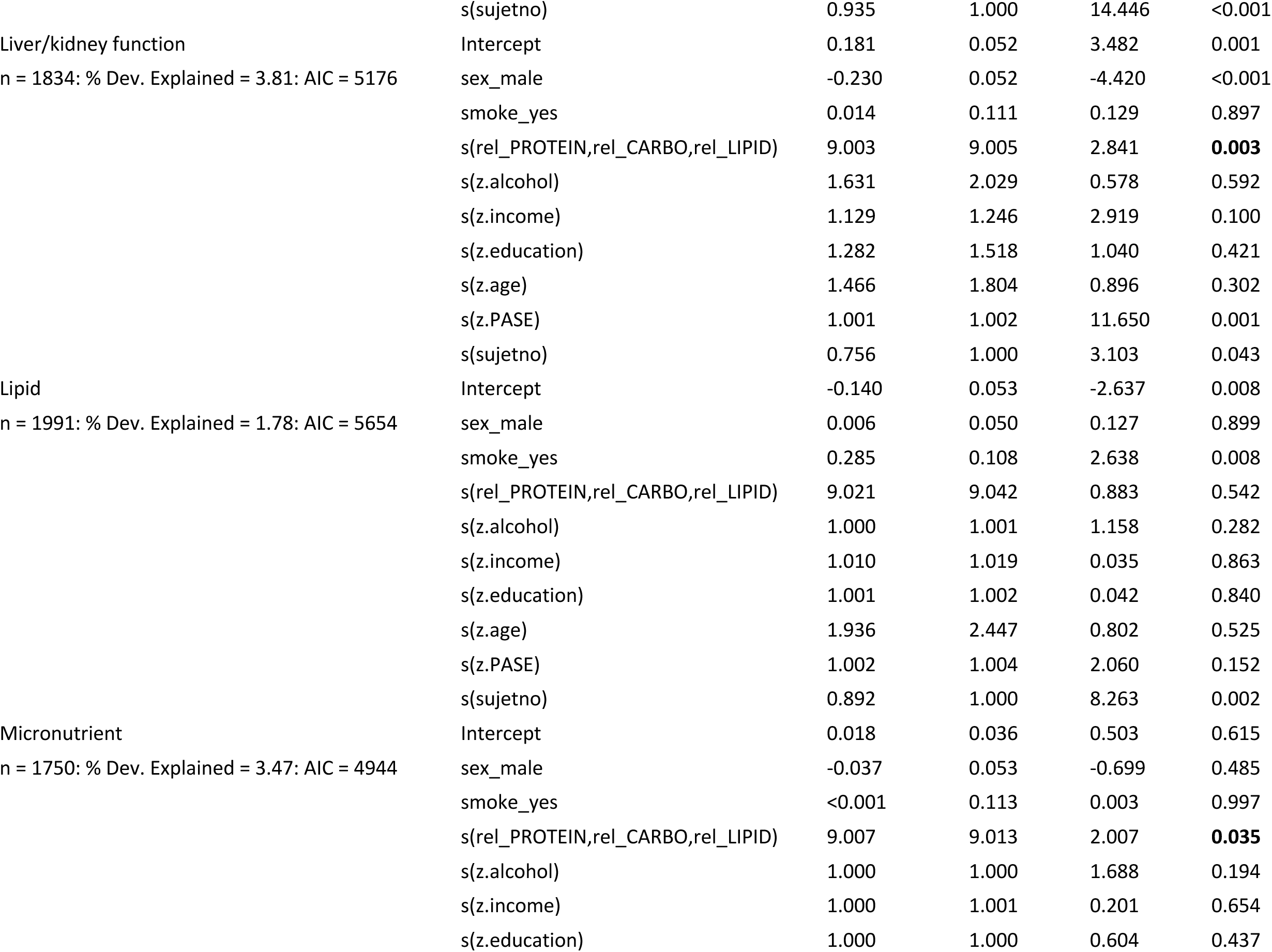
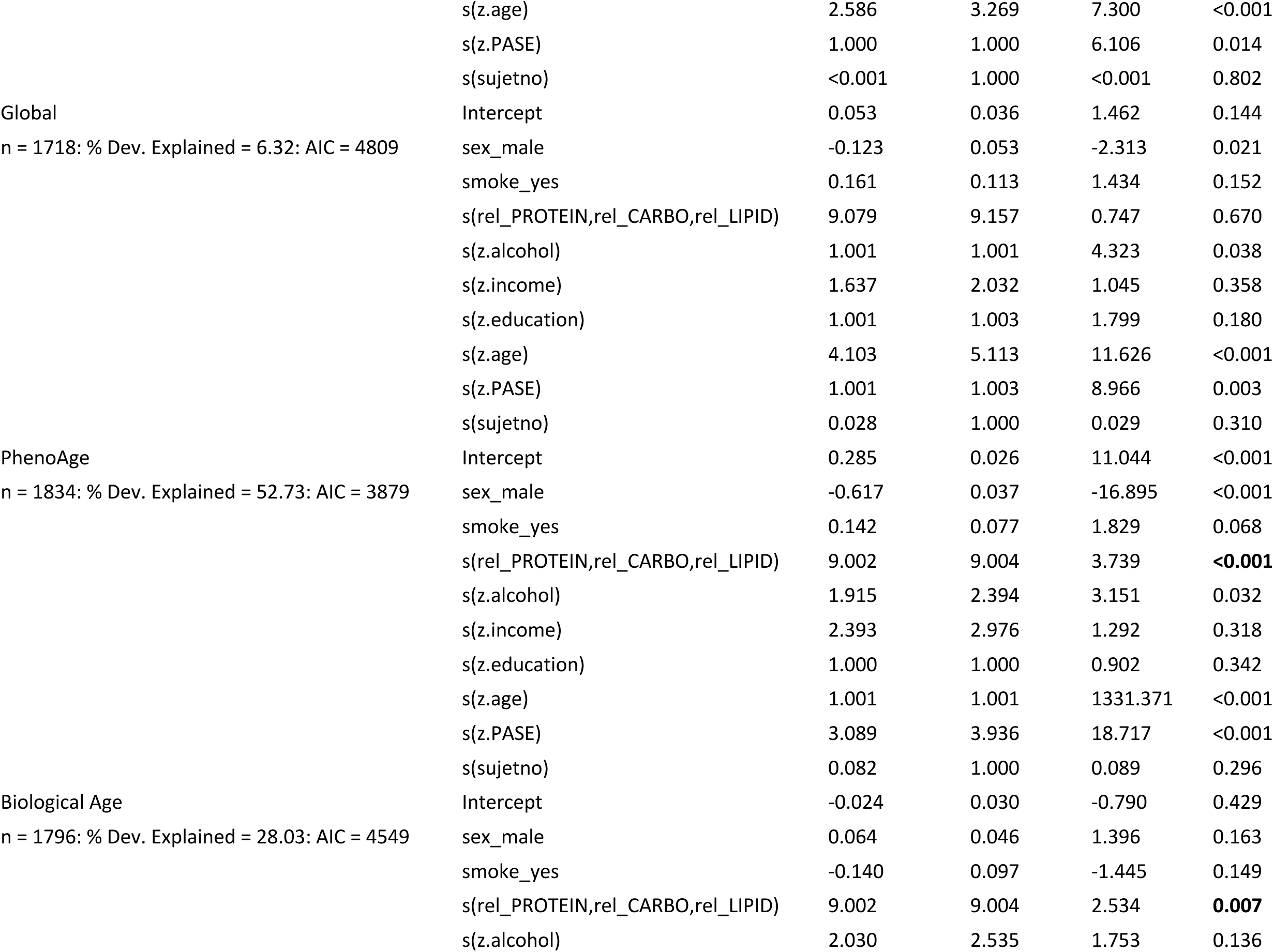
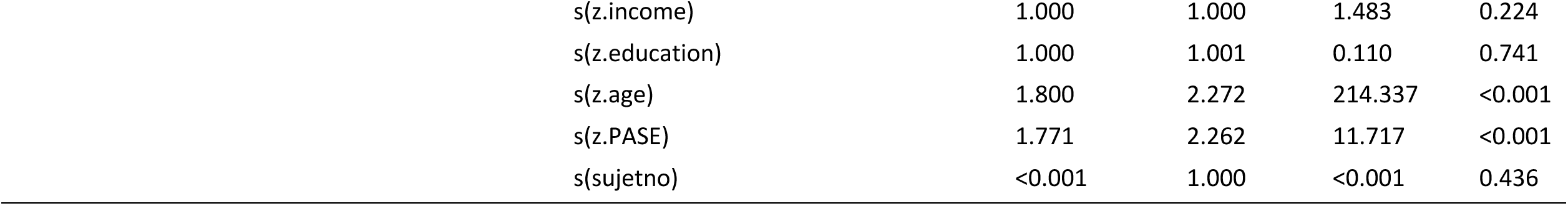
Output of model 3 applied to the inclusive data set with missing income data excluded. For parametric terms estimates, standard error (SE), t and p-values are given. For smooth terms, s(…), estimated degrees of freedom (edf), reference degrees of freedom (Ref. df), F and p-values are given. The sample size (n) is the number of observations, and the random effect for subject ID was fitted as the smooth term s(sujetno). The p-values of significant smooth terms for macronutrients are highlighted in bold.

**Table S5:**
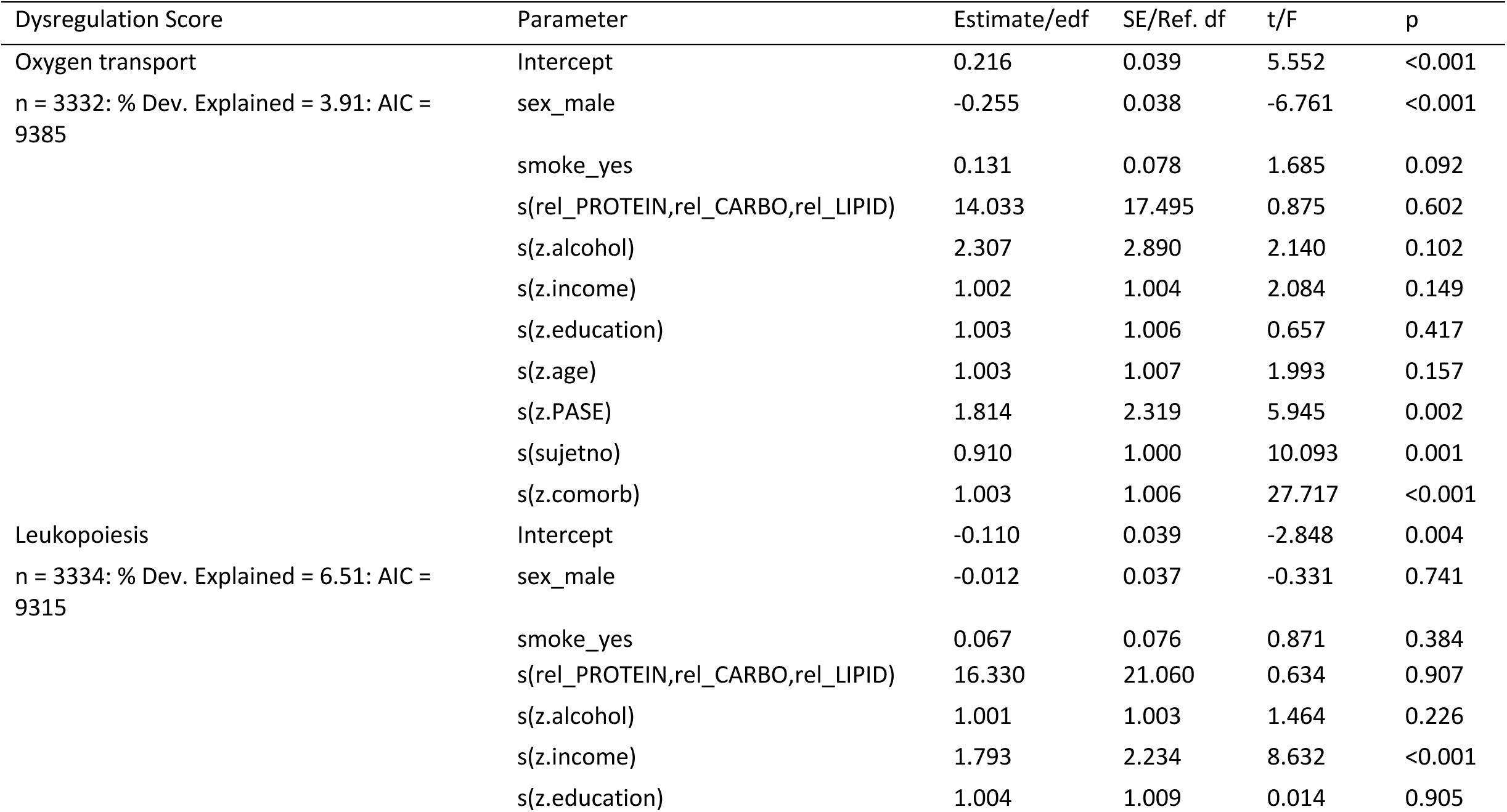
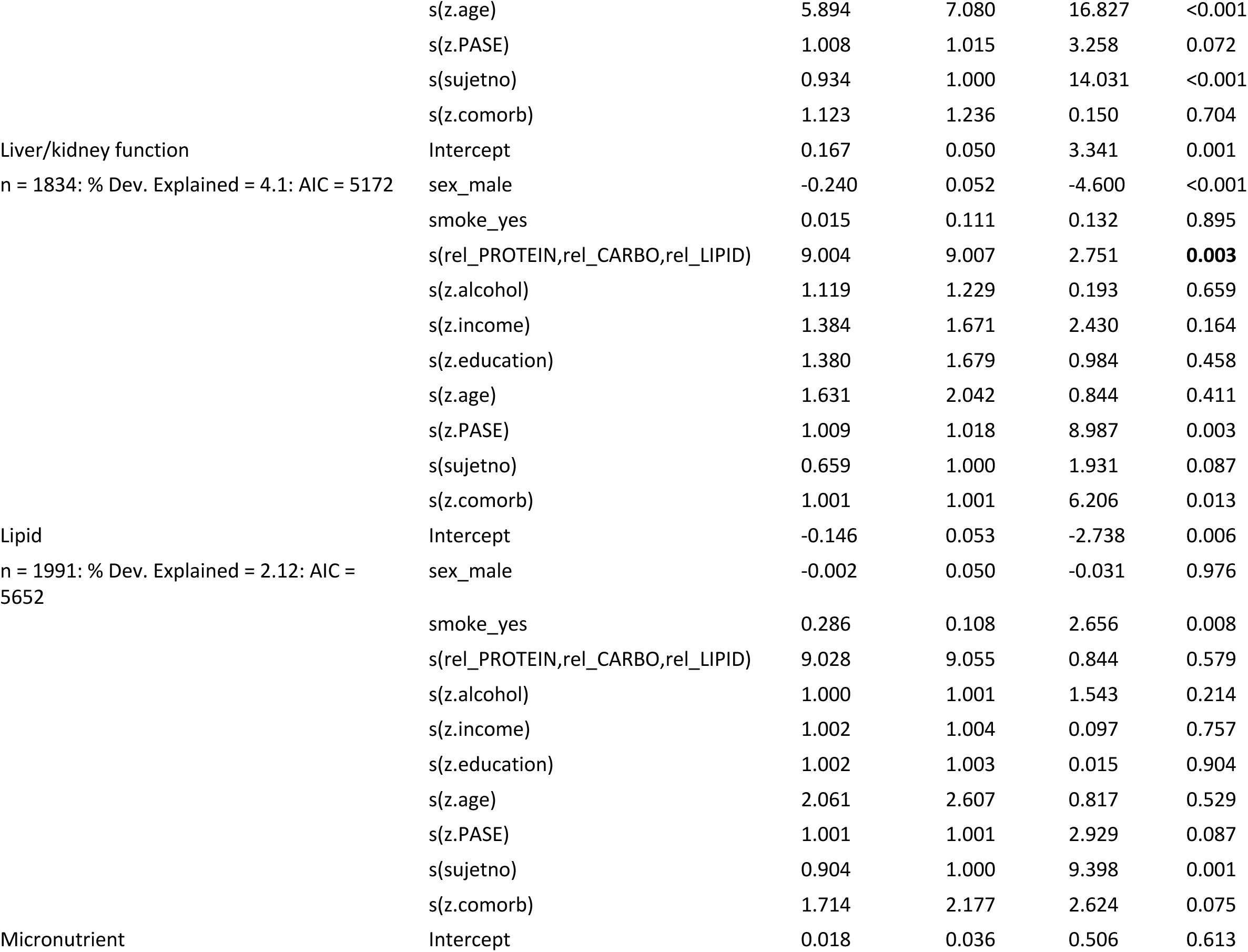
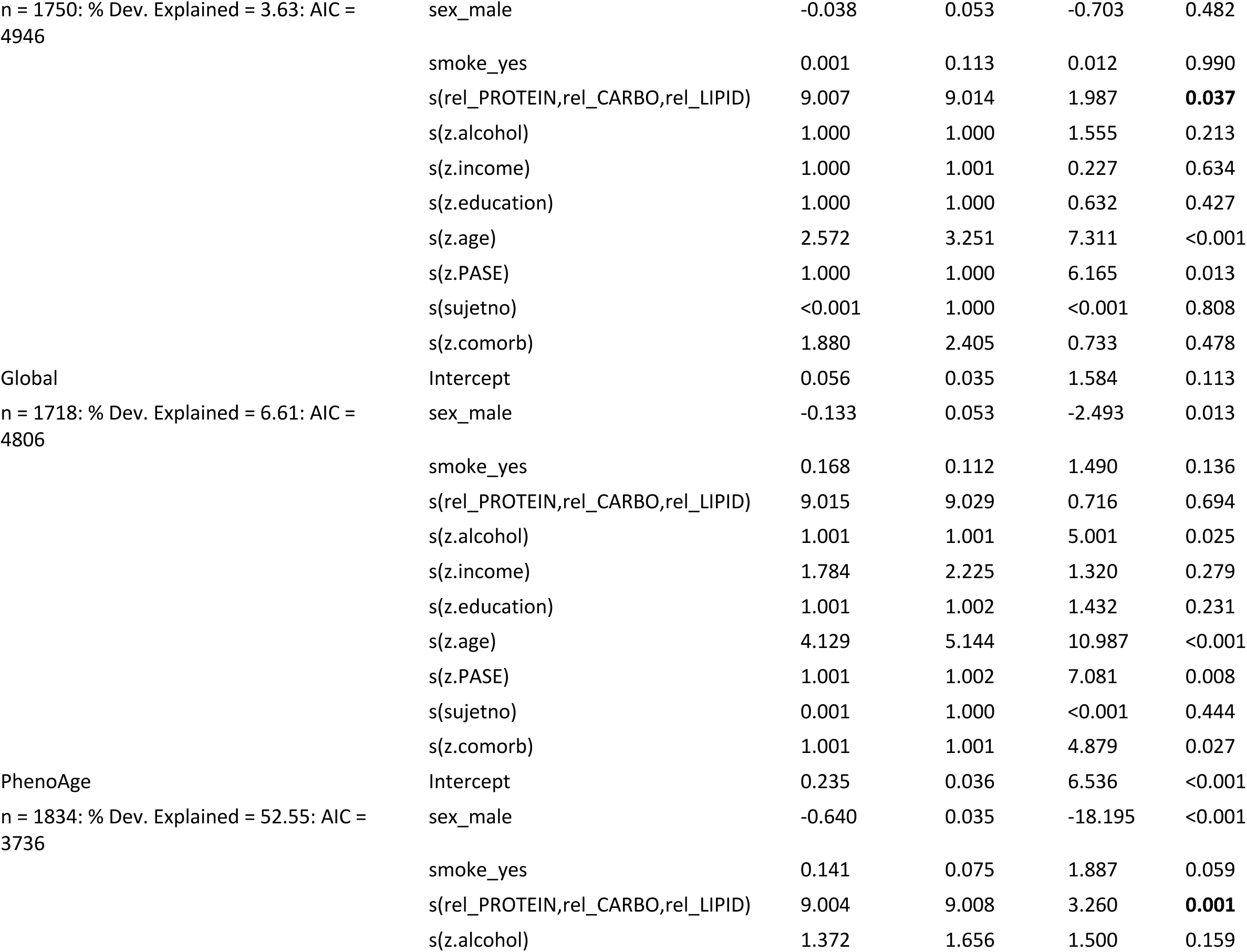
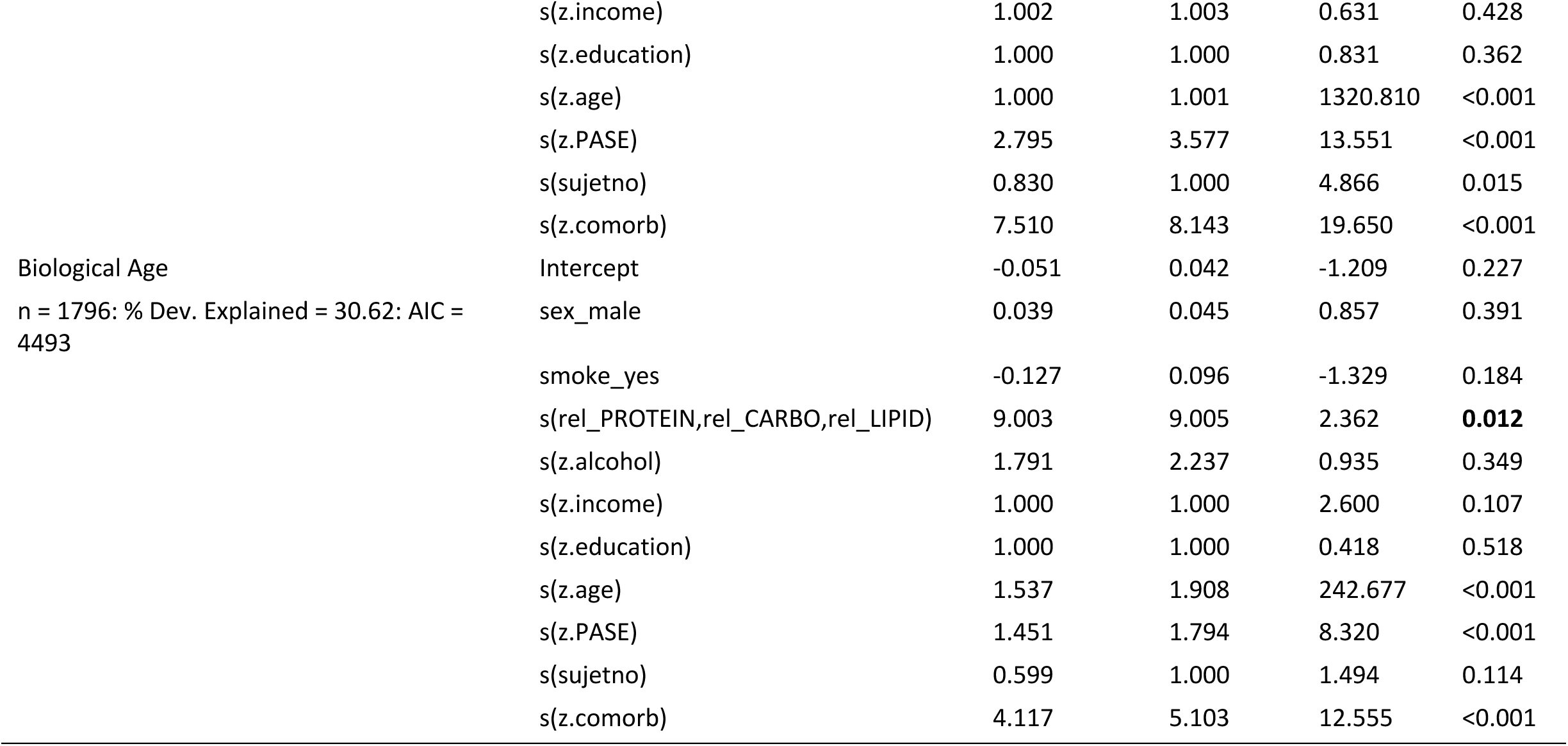
Output of model 4 applied to the inclusive data set with missing income data excluded. For parametric terms estimates, standard error (SE), t and p-values are given. For smooth terms, s(…), estimated degrees of freedom (edf), reference degrees of freedom (Ref. df), F and p-values are given. The sample size (n) is the number of observations, and the random effect for subject ID was fitted as the smooth term s(sujetno). The p-values of significant smooth terms for macronutrients are highlighted in bold.

**Table S6:** Variance statistics from principal component analysis (PCA) of the 7 clusters of micronutrients with highly correlated intakes.

| Cluster | Variance | PC1 | PC2 | PC3 |
| --- | --- | --- | --- | --- |
| 1 | Standard deviation | 497.58 | 160.48 |  |
|  | Proportion of Variance | 0.91 | 0.09 |  |
|  | Cumulative Proportion | 0.91 | 1 |  |
| 2 | Standard deviation | 4.46 | 0.4 |  |
|  | Proportion of Variance | 0.99 | 0.01 |  |
|  | Cumulative Proportion | 0.99 | 1 |  |
| 3 | Standard deviation | 876.56 | 7.43 | 0.28 |
|  | Proportion of Variance | 1 | 0 | 0 |
|  | Cumulative Proportion | 1 | 1 | 1 |
| 4 | Standard deviation | 1329.24 | 3.37 |  |
|  | Proportion of Variance | 1 | 0 |  |
|  | Cumulative Proportion | 1 | 1 |  |
| 5 | Standard deviation | 1752.73 | 45.51 |  |
|  | Proportion of Variance | 1 | 0 |  |
|  | Cumulative Proportion | 1 | 1 |  |
| 6 | Standard deviation | 1.84 | 0.38 |  |
|  | Proportion of Variance | 0.96 | 0.04 |  |
|  | Cumulative Proportion | 0.96 | 1 |  |
| 7 | Standard deviation | 13.59 | 4.58 |  |
|  | Proportion of Variance | 0.9 | 0.1 |  |
|  | Cumulative Proportion | 0.9 | 1 |  |

**Table S7:** Numbers of models with significant 3-way smooth terms for effects of micronutrients on different combinations of outcomes (micronutrient-specific models); 1 indicates the outcome score is in the group (i.e. 1, 0, 0, 0, 0, 0 indicates leukopoiesis dysregulation alone, where 1, 1, 1, 1, 1, 1 indicates all scores).

| Leukopoiesis | Liver/kidney<br>function | Micronutrient | Global | PhenoAge | Biological<br>Age | Number of<br>Models |
| --- | --- | --- | --- | --- | --- | --- |
| 1 | 0 | 0 | 0 | 0 | 0 | 0 |
| 0 | 1 | 0 | 0 | 0 | 0 | 18 |
| 1 | 1 | 0 | 0 | 0 | 0 | 0 |
| 0 | 0 | 1 | 0 | 0 | 0 | 0 |
| 1 | 0 | 1 | 0 | 0 | 0 | 0 |
| 0 | 1 | 1 | 0 | 0 | 0 | 0 |
| 1 | 1 | 1 | 0 | 0 | 0 | 0 |
| 0 | 0 | 0 | 1 | 0 | 0 | 4 |
| 1 | 0 | 0 | 1 | 0 | 0 | 3 |
| 0 | 1 | 0 | 1 | 0 | 0 | 2 |
| 1 | 1 | 0 | 1 | 0 | 0 | 0 |
| 0 | 0 | 1 | 1 | 0 | 0 | 0 |
| 1 | 0 | 1 | 1 | 0 | 0 | 0 |
| 0 | 1 | 1 | 1 | 0 | 0 | 0 |
| 1 | 1 | 1 | 1 | 0 | 0 | 0 |
| 0 | 0 | 0 | 0 | 1 | 0 | 28 |
| 1 | 0 | 0 | 0 | 1 | 0 | 0 |
| 0 | 1 | 0 | 0 | 1 | 0 | 15 |
| 1 | 1 | 0 | 0 | 1 | 0 | 0 |
| 0 | 0 | 1 | 0 | 1 | 0 | 3 |
| 1 | 0 | 1 | 0 | 1 | 0 | 0 |
| 0 | 1 | 1 | 0 | 1 | 0 | 1 |
| 1 | 1 | 1 | 0 | 1 | 0 | 0 |
| 0 | 0 | 0 | 1 | 1 | 0 | 0 |
| 1 | 0 | 0 | 1 | 1 | 0 | 2 |
| 0 | 1 | 0 | 1 | 1 | 0 | 1 |
| 1 | 1 | 0 | 1 | 1 | 0 | 0 |
| 0 | 0 | 1 | 1 | 1 | 0 | 0 |
| 1 | 0 | 1 | 1 | 1 | 0 | 0 |
| 0 | 1 | 1 | 1 | 1 | 0 | 0 |
| 1 | 1 | 1 | 1 | 1 | 0 | 0 |
| 0 | 0 | 0 | 0 | 0 | 1 | 84 |
| 1 | 0 | 0 | 0 | 0 | 1 | 1 |
| 0 | 1 | 0 | 0 | 0 | 1 | 19 |
| 1 | 1 | 0 | 0 | 0 | 1 | 0 |
| 0 | 0 | 1 | 0 | 0 | 1 | 20 |
| 1 | 0 | 1 | 0 | 0 | 1 | 0 |
| 0 | 1 | 1 | 0 | 0 | 1 | 4 |
| 1 | 1 | 1 | 0 | 0 | 1 | 0 |
| 0 | 0 | 0 | 1 | 0 | 1 | 3 |
| 1 | 0 | 0 | 1 | 0 | 1 | 1 |
| 0 | 1 | 0 | 1 | 0 | 1 | 1 |
| 1 | 1 | 0 | 1 | 0 | 1 | 0 |
| 0 | 0 | 1 | 1 | 0 | 1 | 0 |
| 1 | 0 | 1 | 1 | 0 | 1 | 0 |
| 0 | 1 | 1 | 1 | 0 | 1 | 0 |
| 1 | 1 | 1 | 1 | 0 | 1 | 0 |
| 0 | 0 | 0 | 0 | 1 | 1 | 251 |
| 1 | 0 | 0 | 0 | 1 | 1 | 10 |
| 0 | 1 | 0 | 0 | 1 | 1 | 113 |
| 1 | 1 | 0 | 0 | 1 | 1 | 1 |
| 0 | 0 | 1 | 0 | 1 | 1 | 112 |
| 1 | 0 | 1 | 0 | 1 | 1 | 2 |
| 0 | 1 | 1 | 0 | 1 | 1 | 40 |
| 1 | 1 | 1 | 0 | 1 | 1 | 0 |
| 0 | 0 | 0 | 1 | 1 | 1 | 6 |
| 1 | 0 | 0 | 1 | 1 | 1 | 84 |
| 0 | 1 | 0 | 1 | 1 | 1 | 6 |
| 1 | 1 | 0 | 1 | 1 | 1 | 4 |
| 0 | 0 | 1 | 1 | 1 | 1 | 1 |
| 1 | 0 | 1 | 1 | 1 | 1 | 48 |
| 0 | 1 | 1 | 1 | 1 | 1 | 0 |
| 1 | 1 | 1 | 1 | 1 | 1 | 17 |

**Table S8:**
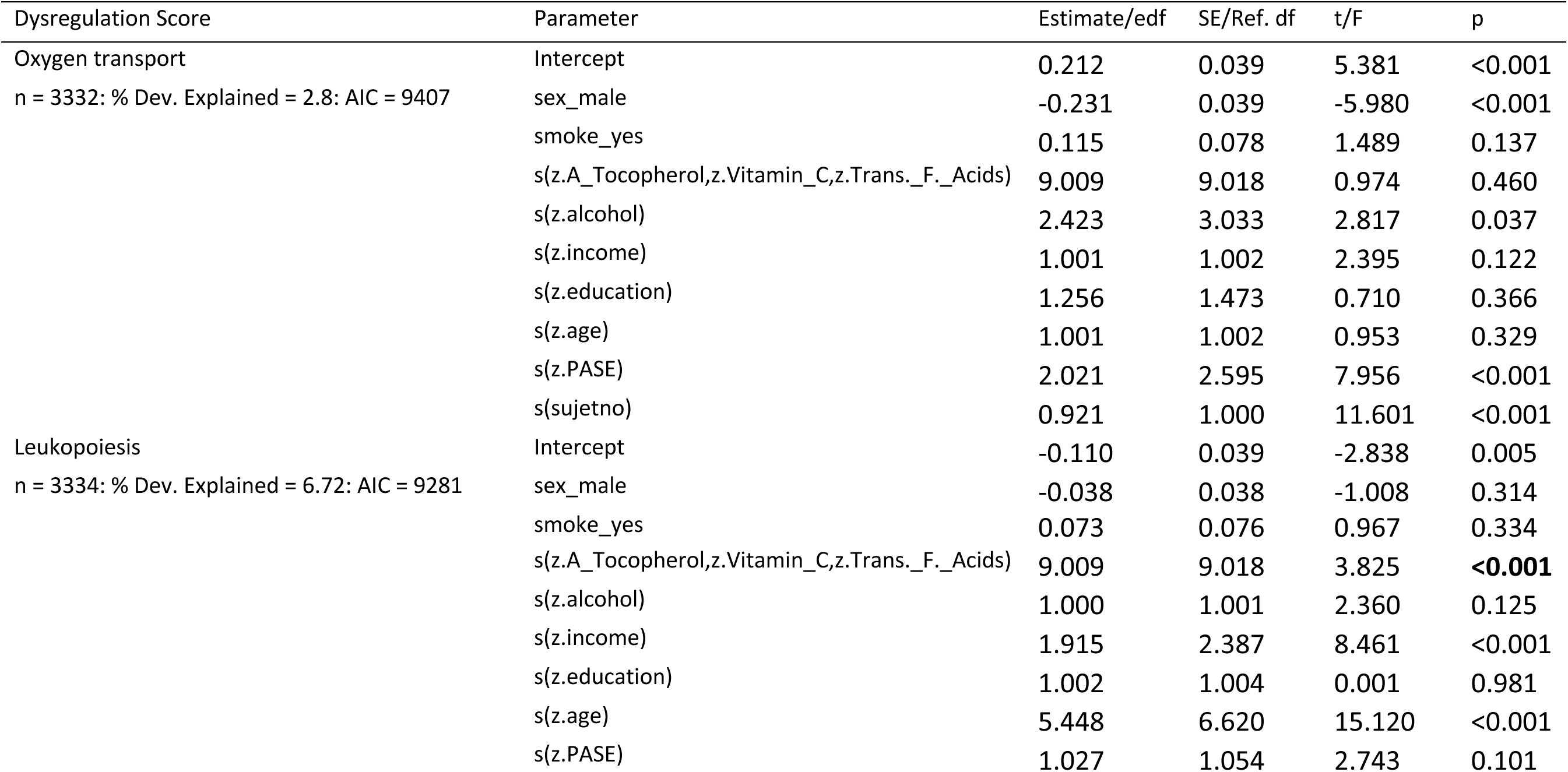
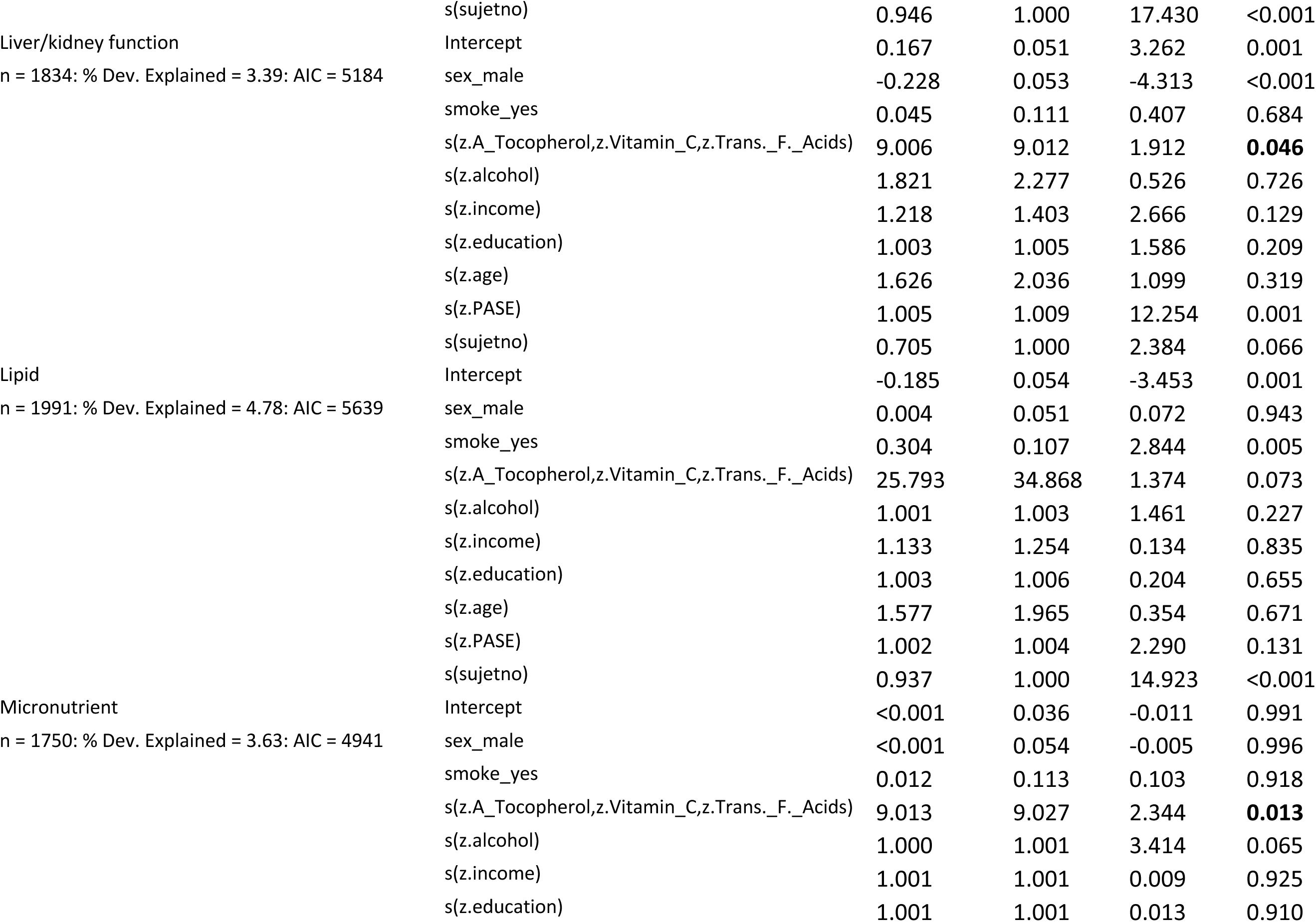
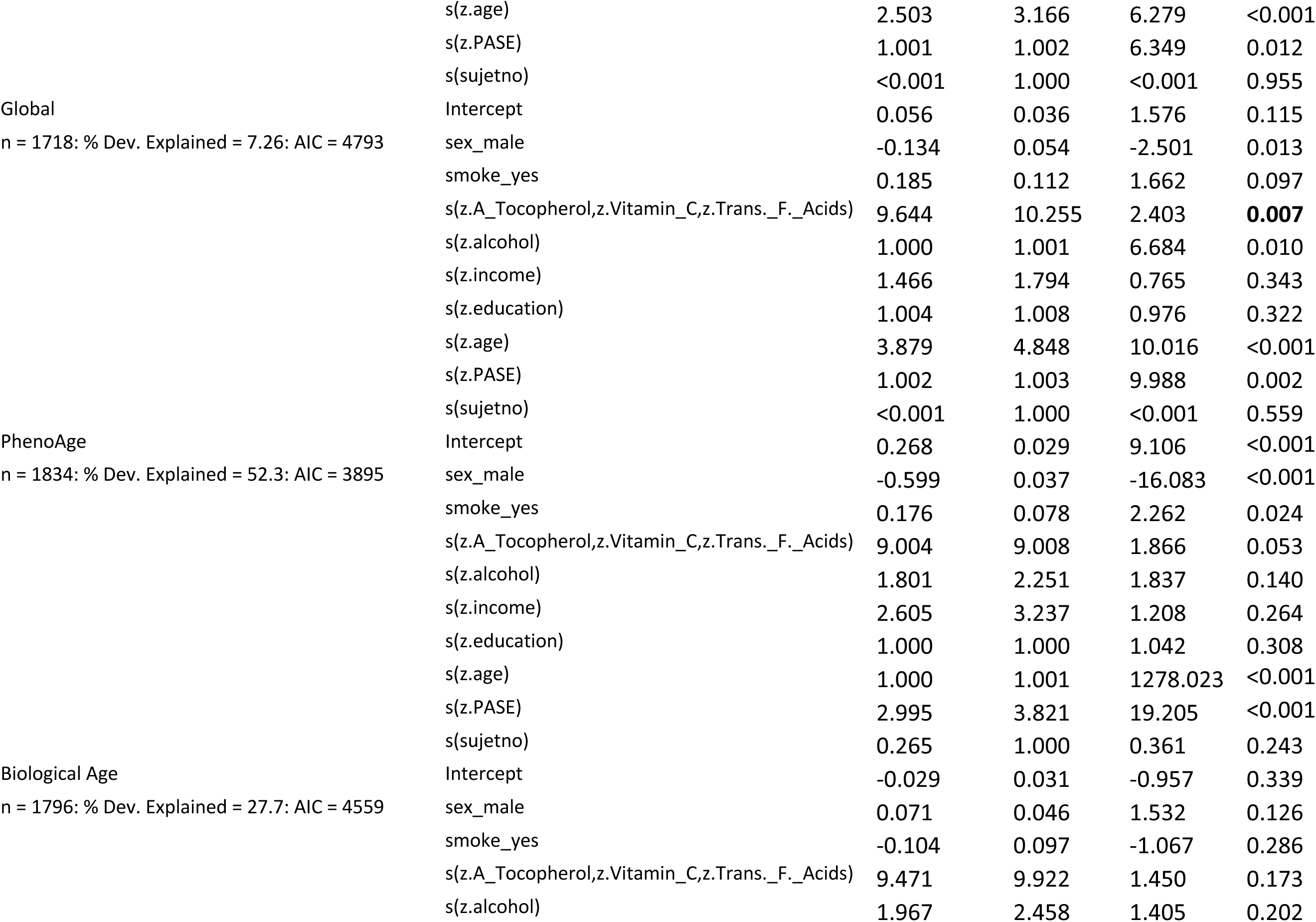
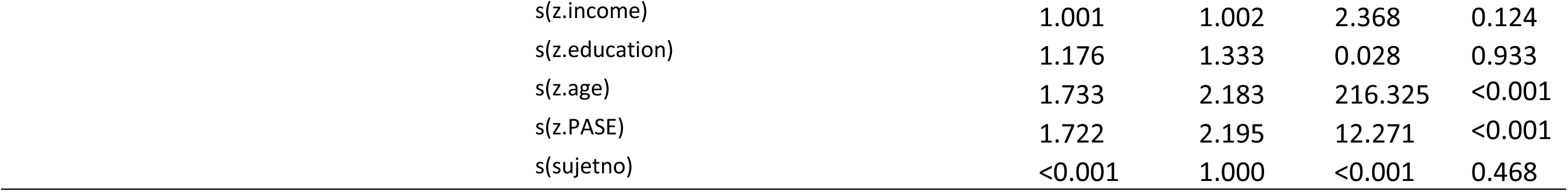
Output of model 5 applied to the inclusive data set with missing income data excluded. For parametric terms estimates, standard error (SE), t and p-values are given. For smooth terms, s(…), estimated degrees of freedom (edf), reference degrees of freedom (Ref. df), F and p-values are given. The sample size (n) is the number of observations, and the random effect for subject ID was fitted as the smooth term s(sujetno). The p-values of significant smooth terms for micronutrients are highlighted in bold.

**Table S9:**
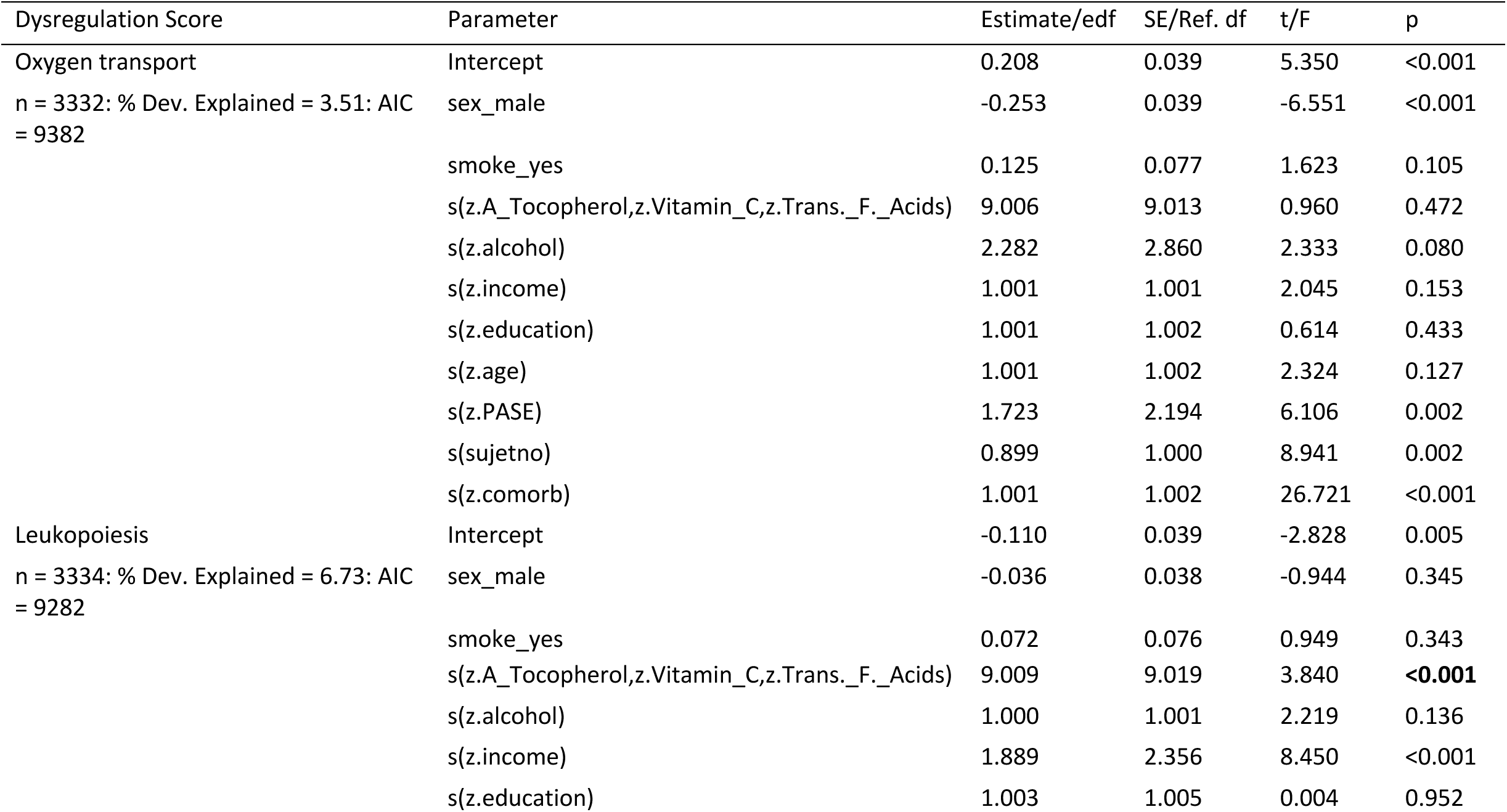
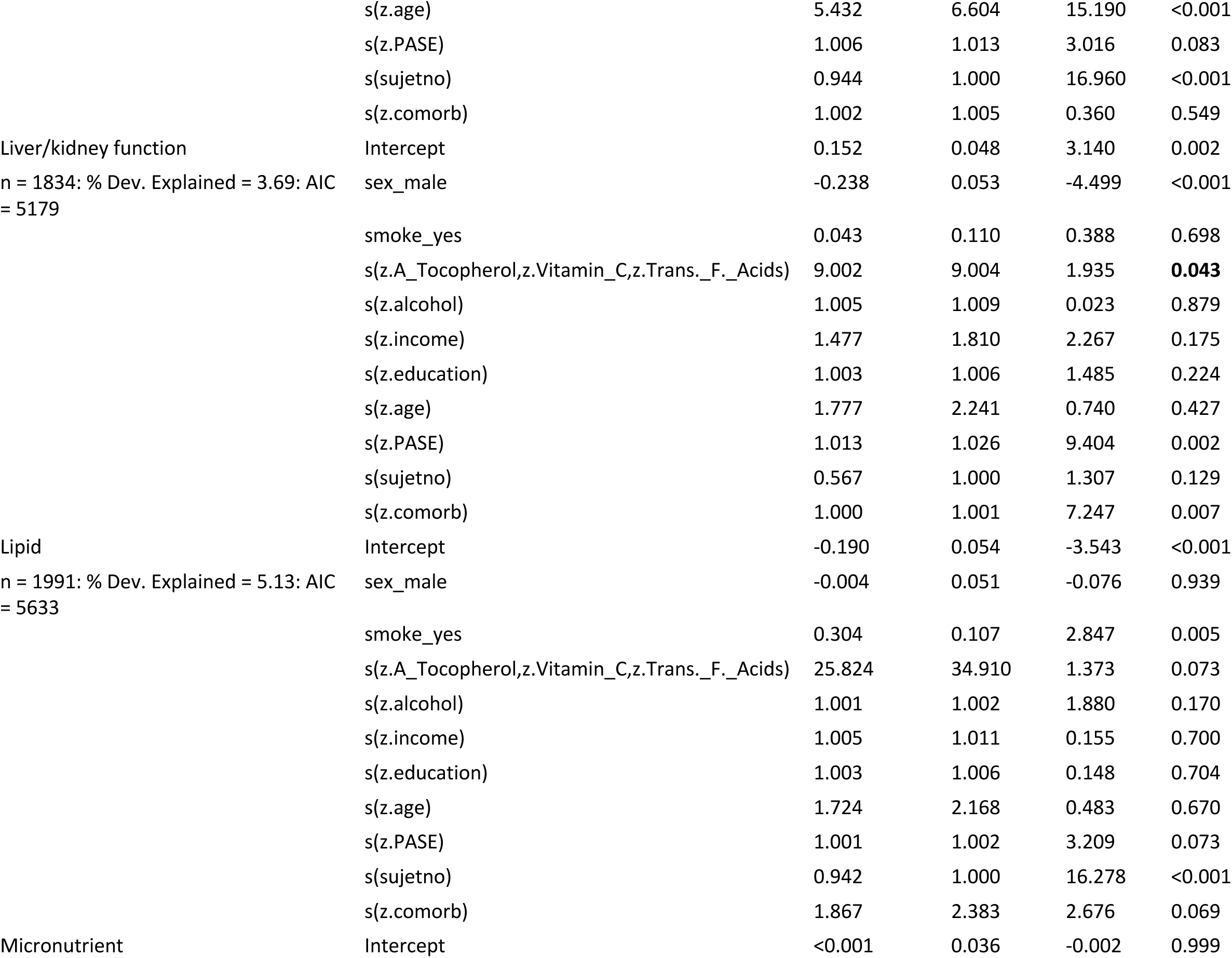
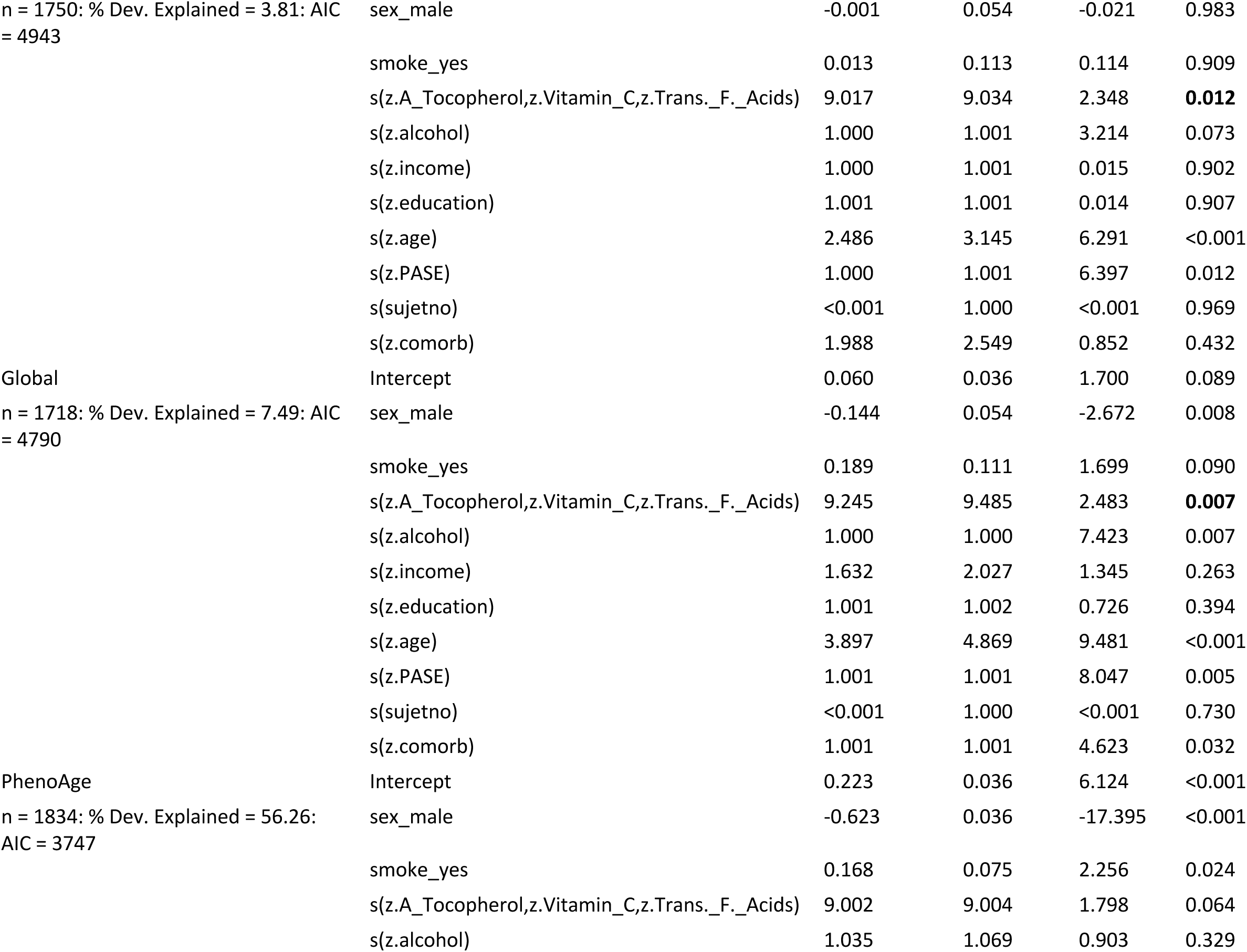
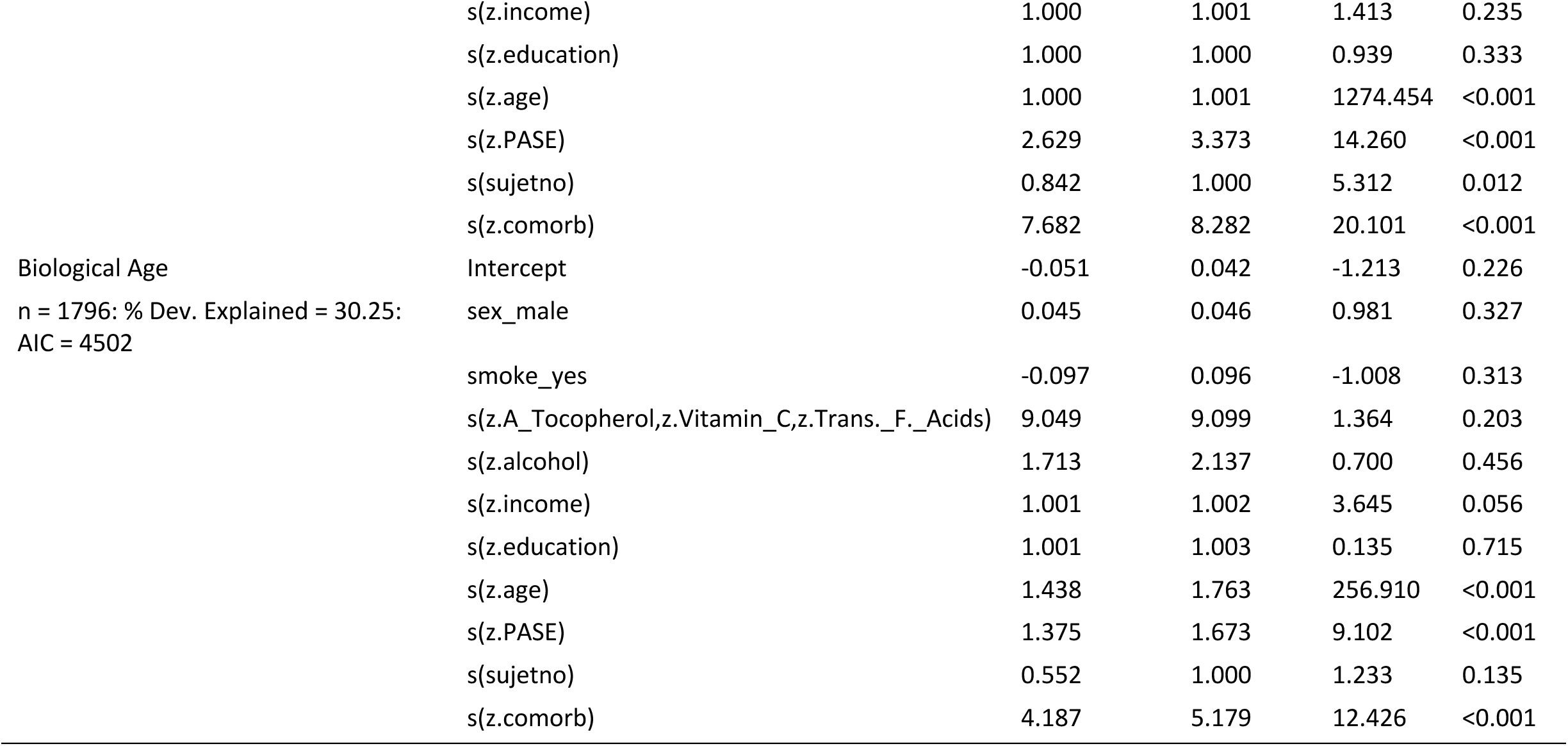
Output of model 6 applied to the inclusive data set with missing income data excluded. For parametric terms estimates, standard error (SE), t and p-values are given. For smooth terms, s(…), estimated degrees of freedom (edf), reference degrees of freedom (Ref. df), F and p-values are given. The sample size (n) is the number of observations, and the random effect for subject ID was fitted as the smooth term s(sujetno). The p-values of significant smooth terms for micronutrients are highlighted in bold.

**Table S10:**
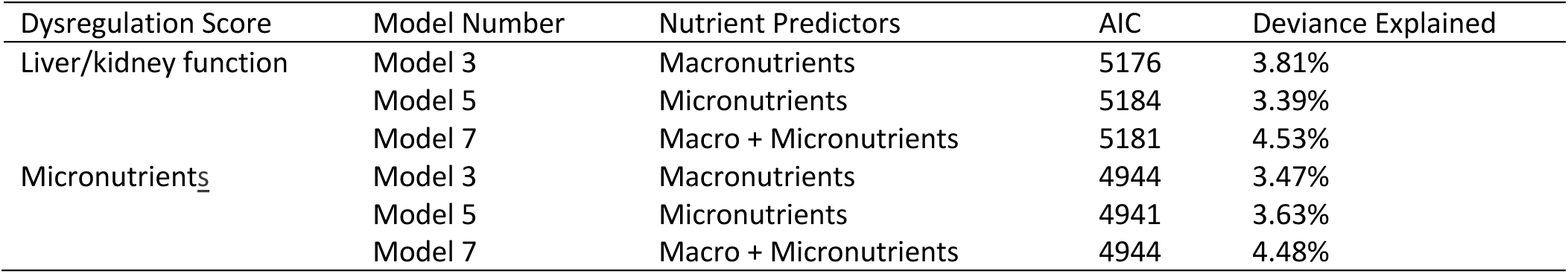
Akaike information criterion (AIC) and deviance explained (%) by models 3, 5, and 7, applied to the inclusive data set with missing income data excluded, for liver/kidney and micronutrients dysregulation. These models contained, alcohol intake, sex, smoking status, income, education level, age, and physical activity (PASE) alongside the nutritional predictors stated.

| Dysregulation Score | Model Number | Nutrient Predictors | AIC | Deviance Explained |
| --- | --- | --- | --- | --- |
| Liver/kidney function | Model 3 | Macronutrients | 5176 | 3.81% |
|  | Model 5 | Micronutrients | 5184 | 3.39% |
|  | Model 7 | Macro + Micronutrients | 5181 | 4.53% |
| Micronutrients | Model 3 | Macronutrients | 4944 | 3.47% |
|  | Model 5 | Micronutrients | 4941 | 3.63% |
|  | Model 7 | Macro + Micronutrients | 4944 | 4.48% |

**Table S11.** Akaike information criterion (AIC) for mixture models (MM) (see 1) of micronutrient intake (*Z*-transformed) as a function of dietary macronutrient composition (% total energy from protein, carbohydrates and fats; see text S5), as applied to the inclusive data set with missing income data excluded. The models favoured by AIC for each macronutrient is highlighted in bold italics.

| Micronutrient | MM 1 (null) | MM 2 | MM 3 | MM 4 | MM 5 |
| --- | --- | --- | --- | --- | --- |
| Cluster 1 PC1 | 10131 | 10057 | 10044 | 10040 | ***10036*** |
| Cluster 2 PC1 | 10131 | 10120 | ***10096*** | 10098 | 10098 |
| Magnesium | 10131 | 10079 | ***10068*** | 10070 | 10070 |
| Cluster 3 PC1 | 10131 | 10012 | ***10000*** | 10005 | 10001 |
| Sodium | 10131 | 10008 | 9996 | 9987 | ***9985*** |
| Zinc | 10131 | 9890 | ***9870*** | 9875 | 9872 |
| Selenium | 10131 | 9870 | 9862 | 9863 | ***9858*** |
| Cluster 4 PC1 | 10131 | ***10116*** | 10119 | 10123 | 10119 |
| A Tocopherol | 10131 | 10004 | ***10001*** | 10004 | 10000 |
| Lycopene | 10131 | ***10123*** | 10123 | 10129 | 10125 |
| Cluster 5 PC1 | 10131 | ***10088*** | 10088 | 10090 | 10089 |
| Vitamin D | 10131 | ***10067*** | 10069 | 10071 | 10068 |
| Vitamin C | 10131 | 9949 | ***9943*** | 9943 | 9944 |
| Cluster 6 PC1 | 10131 | 10047 | ***10025*** | 10031 | 10027 |
| Folic Acid Equiv. | 10131 | 10090 | ***10084*** | 10087 | 10085 |
| Cholesterol | 10131 | 9429 | 9417 | ***9414*** | 9417 |
| Trans. F. Acids | 10131 | ***9848*** | 9850 | 9855 | 9852 |
| Cluster 7 PC1 | 10131 | 8063 | 8059 | ***8056*** | 8057 |
| Poly. F. Acids | 10131 | 9163 | 9157 | ***9155*** | 9155 |

## Supplementary Figures

**Figure S1.**
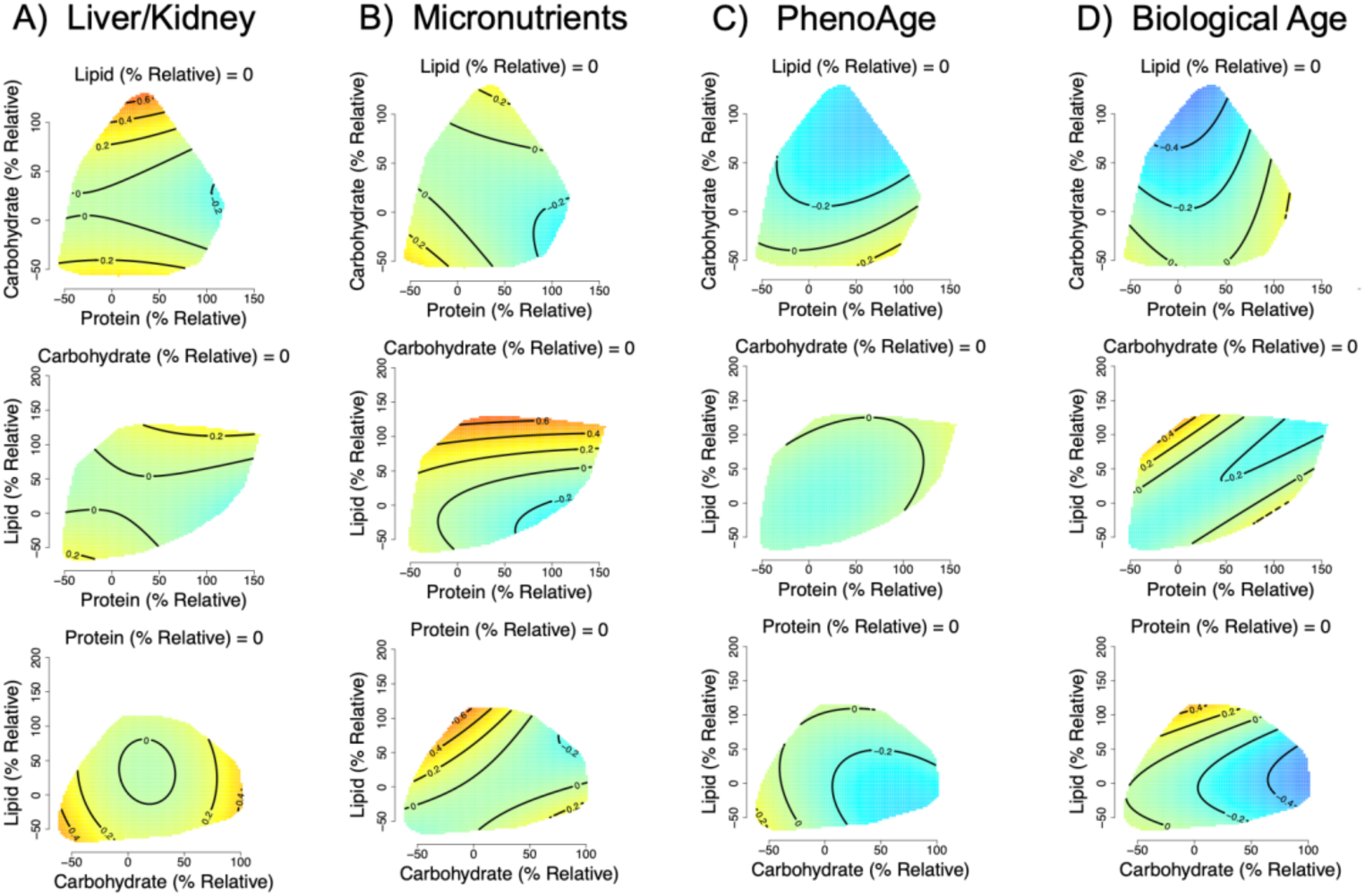
Effects of relative dietary macronutrient intake (relative to the required intake based on age, weight, height sex and physical activity level; see text S1) on A) liver/kidney function dysregulation, (GAM three-way smooth term: edf=9, Ref. df=9, F=4, p<0.001, Dev. Expl.=1.5%), B) micronutrient dysregulation, (GAM three-way smooth term: edf=9.2, Ref. df=9.4, F=2.4, p=0.01, Dev. Expl.=0.9%), C) PhenoAge (GAM three-way smooth term: edf=9, Ref. df=9, F=2.8, p<0.01, Dev. Expl.=1.6%) and D) biological age (GAM three-way smooth term: edf=9, Ref. df=9, F=2.9, p<0.01, Dev. Expl.=1.8%) score as predicted by model 2 applied to the inclusive dataset with imputation of missing income data. Surfaces across the top row show effects of protein (x-axis), and carbohydrate (y-axis) intake, those across the middle row protein and lipid, and the bottom row is carbohydrate and lipid. The third macronutrient is held at the values given on all panels. Warm colours indicate high dysregulation, and cool colours low dysregulation. All scores were Z-transformed to one SD, and surfaces colours are scaled such that deep blue and red represent effects of at least -0.8 and 0.8 (conventionally considered an effect of large biological magnitude (6)). Individuals with a relative intake value of 100, eat 100% more of that macronutrient per day (in kJ) than is predicted to be typical for the population given their age, sex, weight, height and level of physical activity level. Conversely, individuals with a relative intake value of 0 would eat the required amount of that macronutrient per day.

**Figure S2.**
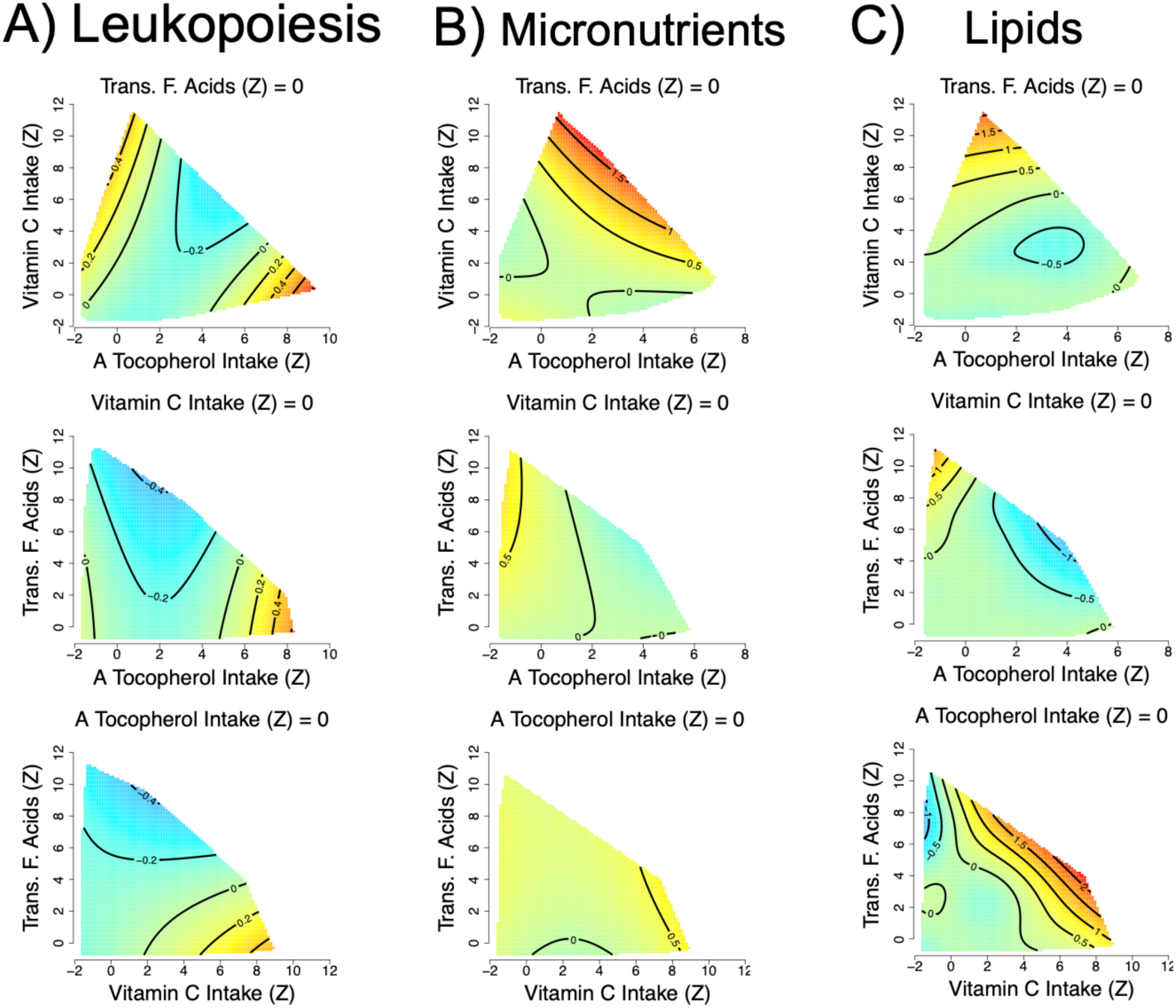
Effects of total dietary micronutrient (*α*-tocopherol, vitamin C and trans-fatty acid intake) intake on A) leukopoiesis (GAM three-way smooth term: edf=9, Ref. df=9, F=2.5, p<0.01, Dev. Expl.=5%), B) micronutrient (GAM three-way smooth term: edf=9, Ref. df=9, F=3.6, p<0.001, Dev. Expl.=4%) and C) lipid (GAM three-way smooth term: edf=28.7, Ref. df=38.4, F=1.5, p<0.05, Dev. Expl.=5%) dysregulation score as predicted by model 6 applied to the inclusive dataset with imputation of missing income data. Intakes have been *Z*-transformed and are thus in units of SD. In all cases predictions assume the micronutrient not displayed on either the x- or y-axis is held at the population mean. Numeric confounding variables included in model 6 were alcohol intake, income, education level, age, physical activity levels (PASE) and number of comorbidities, and predictions assume population mean values. Predictions are for men and assume a non-smoker.

**Figure S3.**
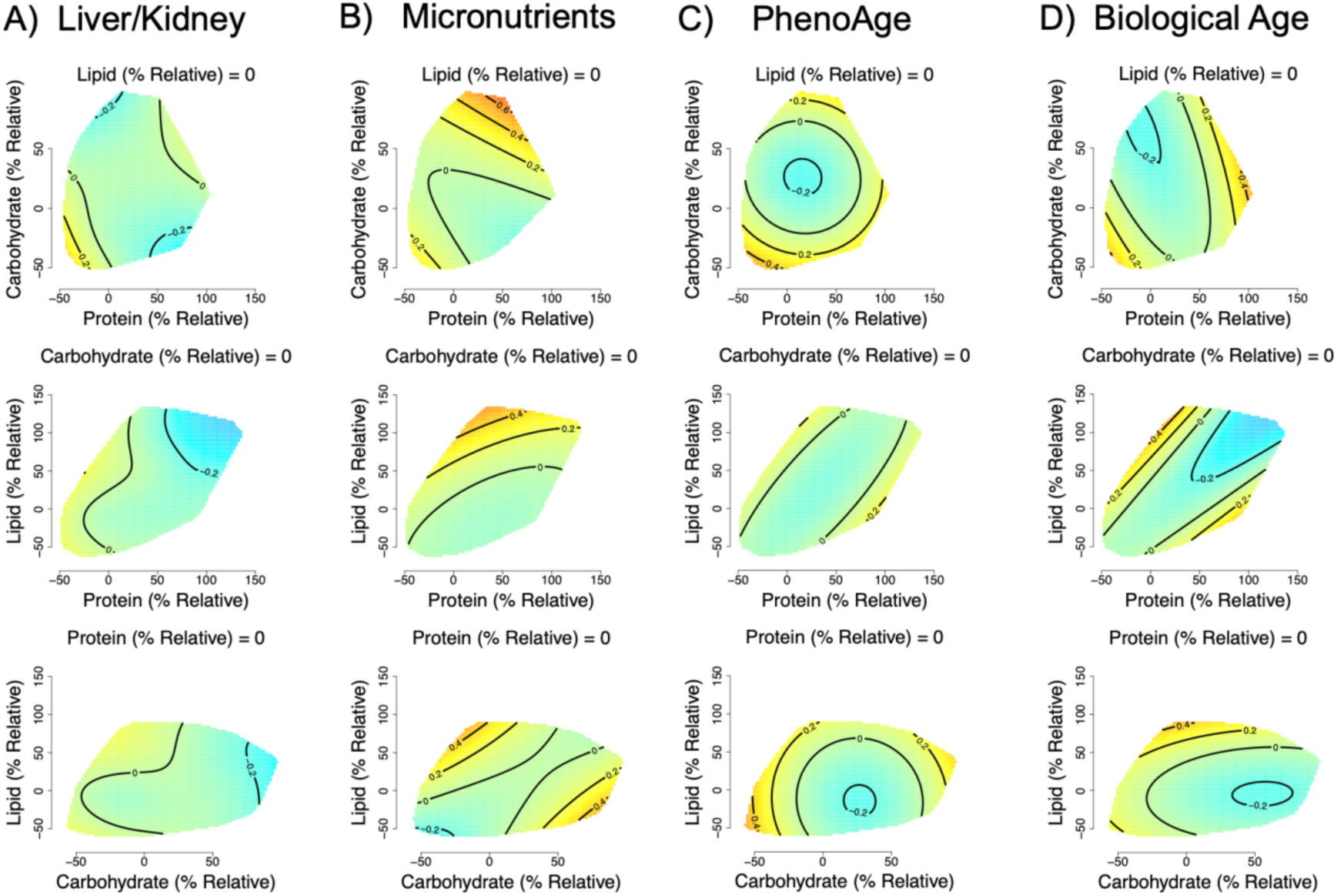
Effects of relative dietary macronutrient intake (relative to typical intake based on age, weight, height sex and physical activity level; see text S1) on A) liver/kidney function (GAM three-way smooth term: edf=13.2, Ref. df=15.9, F=1.1, p=0.34, Dev. Expl.=2.7%) and B) micronutrient (GAM three-way smooth term: edf=9, Ref. df=9, F=0.9, p=0.57, Dev. Expl.=0.9%) dysregulation score as predicted by model 1 applied to the exclusive dataset. Effects of relative dietary macronutrient intake (corrected for age, weight, height sex and PASE) on C) PhenoAge (GAM three-way smooth term: edf=9, Ref. df=9, F=2.6, p=0.005, Dev. Expl.=2.6%) and D) biological age (GAM three-way smooth term: edf=9, Ref. df=9, F=1.7, p=0.08, Dev. Expl.=2%) dysregulation score as predicted by model 2 applied to the exclusive dataset. Surfaces across the top row show effects of protein (x-axis), and carbohydrate (y-axis) intake, those across the middle row protein and lipid, and the bottom row is carbohydrate and lipid. The third macronutrient is held at the values given on all panels. Warm colours indicate high dysregulation, and cool colours low dysregulation. All scores were Z-transformed to one SD, and surfaces colours are scaled such that, values of -0.8 and 0.8 would be red and blue respectively. Individuals with a relative intake value of 100, eat 100% more of that macronutrient per day (in kJ) than is predicted to be typical for the population given their age, sex, weight, height and level of physical activity level. Conversely, individuals with a relative intake value of 0 would eat the required amount of that macronutrient per day.

**Figure S4.**
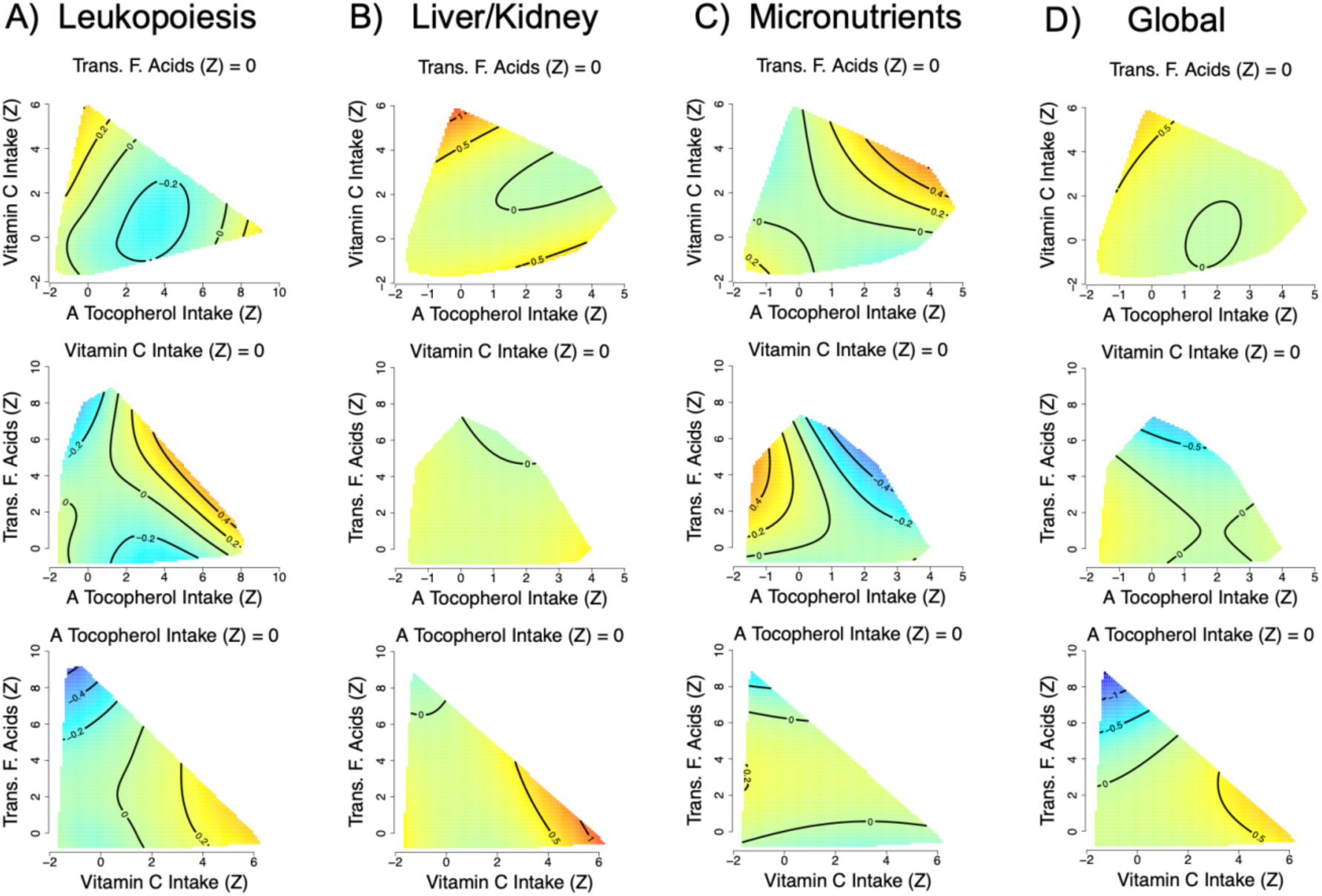
Effects of total dietary micronutrient (*α*-tocopherol, vitamin C and trans-fatty acid intake) intake on A) leukopoiesis (GAM three-way smooth term: edf=12.4, Ref. df=15, F=0.9, p=0.56, Dev. Expl.=8.9%), B) liver/kidney function (GAM three-way smooth term: edf=9, Ref. df=9, F=1, p=0.41, Dev. Expl.=5.25%), C) micronutrient (GAM three-way smooth term: edf=9, Ref. df=9, F=1.03, p=0.41, Dev. Expl.=3.1%) and D) global (GAM three-way smooth term: edf=9, Ref. df=9, F=1.9, p=0.05, Dev. Expl.=8.25%) dysregulation score as predicted by model 6 applied to the exclusive dataset. Intakes have been *Z*-transformed and are thus in units of SD. In all cases predictions assume the micronutrient not displayed on either the x- or y-axis is held at the population mean. Numeric confounding variables included in model 6 were alcohol intake, income, education level, age, physical activity level (PASE) and number of comorbidities, and predictions assume population mean values. Predictions are for men and assume a non-smoker.

**Figure S5.**
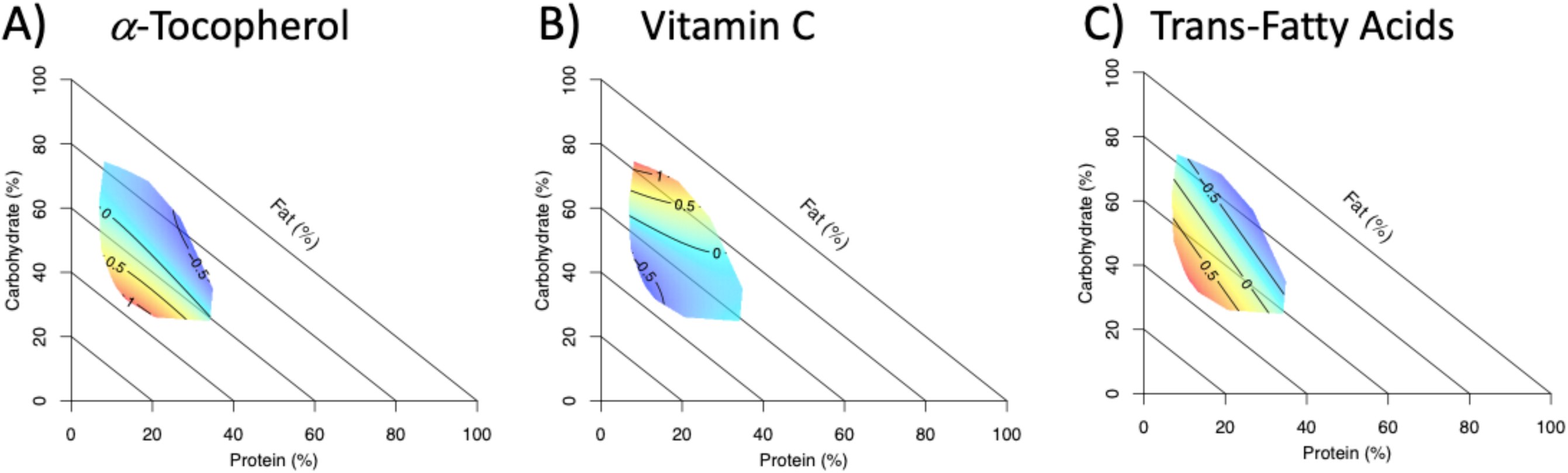
Right angle mixture triangles (5) of associations between dietary macronutrient composition and intake of A) *α*-tocopherol, B) vitamin C and C) trans-fatty acid intake as predicted by AIC favoured mixture models (1, see text S5). Percentage energy from protein and carbohydrates are on the *x*- and *y*-axes respectively, while percentage energy from fats is on the implicit axis. Micronutrient intakes (outcome) have been *z*-transformed such that zero is the population average and units are in one standard deviation.

**Figure S6.**
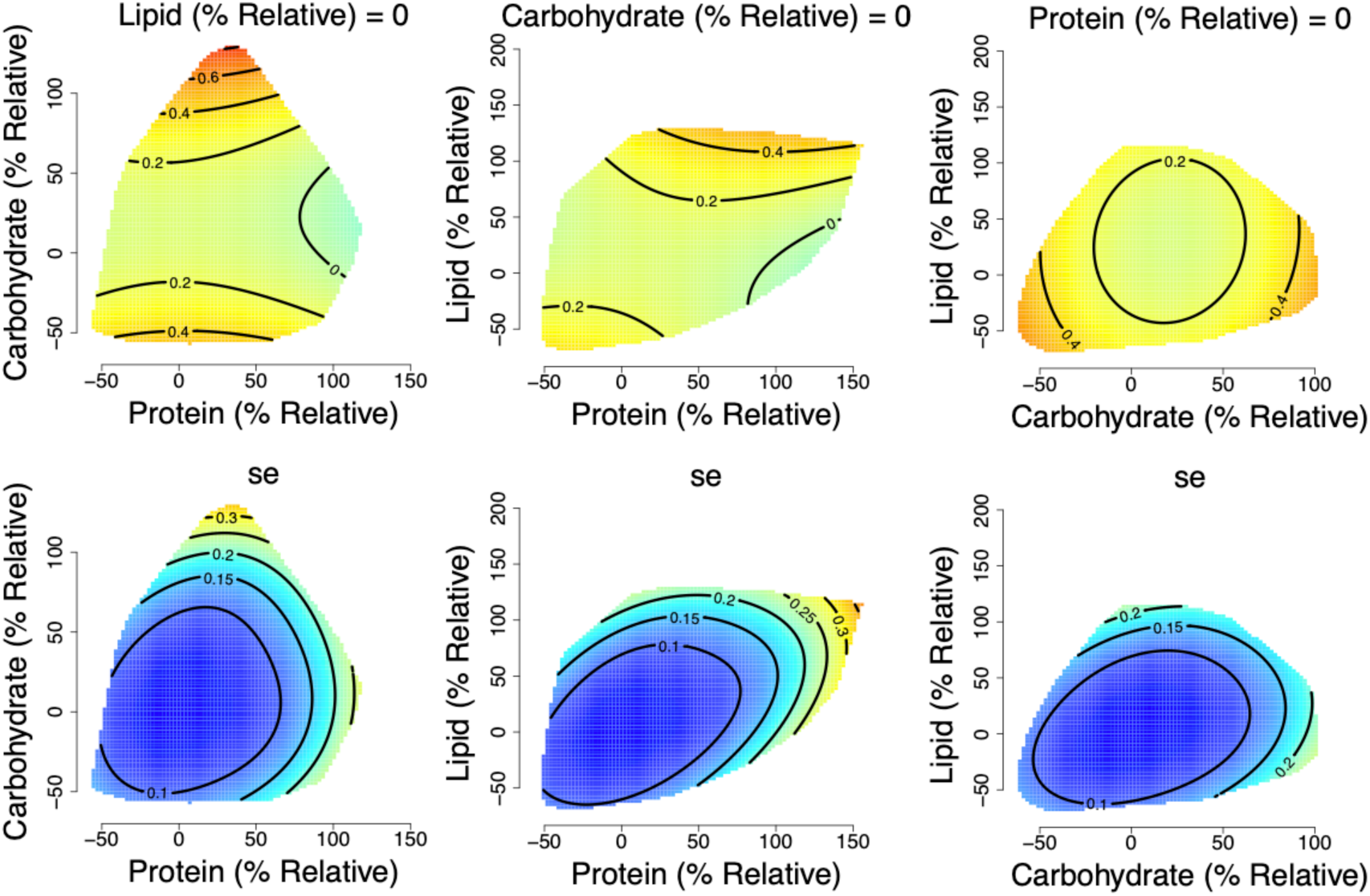
Effects with corresponding standard errors (se) of relative macronutrient intake (relative to estimated requirements; see text S1) on liver/kidney function dysregulation score as predicted by model 4 applied to the inclusive dataset with imputation of missing income data. Left hand surfaces show effects of protein (x-axis), and carbohydrate (y-axis) intake, middle show protein and lipid, and the right is carbohydrate and lipid. The third macronutrient is held at the values given on all panels. Warm colours indicate high dysregulation, and cool colours low dysregulation. All scores were Z-transformed to one SD, and surfaces colours are scaled such that deep blue and red represent effects of at least -0.8 and 0.8 (conventionally considered an effect of large biological magnitude (6)). In all cases predictions assume the macronutrient not displayed on either the x- or y-axis is held at the population median. Numeric confounding variables included in model 4 were alcohol intake, income, education level, age, physical activity levels (PASE) and number of comorbidities, and predictions assume population mean values. Predictions are for men and assume a non-smoker.

